# Genetically and Functionally Distinct Immunoglobulin Heavy Chain Locus Duplication in Bats

**DOI:** 10.1101/2024.08.09.606892

**Authors:** Taylor Pursell, Ashley Reers, Artem Mikelov, Prasanti Kotagiri, James A. Ellison, Christina L. Hutson, Scott D. Boyd, Hannah K. Frank

## Abstract

The genetic locus encoding immunoglobulin heavy chains (IgH) is critical for vertebrate humoral immune responses and diverse antibody repertoires. Immunoglobulin and T cell receptor loci of most bat species have not been annotated, despite the recurrent role of bats as viral reservoirs and sources of zoonotic pathogens. We investigated the genetic structure and function of IgH loci across the largest bat family, Vespertilionidae, focusing on big brown bats *(Eptesicus fuscus*). We discovered that *E. fuscus* and ten other species within Vespertilionidae have two complete, functional, and distinct immunoglobulin heavy chain loci on separate chromosomes. This locus organization is previously unknown in mammals, but is reminiscent of more limited duplicated loci in teleost fish. Single cell transcriptomic data validate functional rearrangement and expression of immunoglobulin heavy chains of both loci in the expressed repertoire of *Eptesicus fuscus*, with maintenance of allelic exclusion, bias of usage toward the smaller and more compact IgH locus, and evidence of differential selection of antigen-experienced B cells and plasma cells varying by IgH locus use. This represents a unique mechanism for mammalian humoral immunity and may contribute to bat resistance to viral pathogenesis.

## Main

Bats are of immunological interest due to their unique relationship with infectious diseases. While bats host more zoonotic viruses per species than any other mammalian order^1^, they often do not experience overt pathology^2^. Multiple adaptations in their innate immunity which impart tolerance to these viruses have been characterized^3–6^, but bat adaptive immunity remains understudied. Bats mount humoral responses to a range of viruses as evidenced by serological studies, and serology is widely used to estimate viral prevalence in bat species across the globe^7–11^, but the genetic and molecular basis for the formation and evolution of bat B cell receptor repertoires and antibody responses is largely uncharacterized.

The genes encoding the heavy and light chains of mammalian immunoglobulin (Ig) proteins are known to be generated via DNA recombination at a single heavy chain locus and one of the two light chain loci in most mammals, designated kappa and lambda. Each of these loci contain arrays of variable (IGHV/IGKV/IGLV), diversity (IGHD), joining (IGHJ/IGKJ/IGLJ), and constant (IGHC/IGKC/IGLC) genes. The number, diversity and organization of these genes contributes to the scale of the resulting immunoglobulin repertoire and varies across species. The phylogenetic order comprising the bats, Chiroptera, contains over 1,400 species with diverse biology and modes of living, and is second only to rodents in mammalian species richness. Despite this diversity, studies characterizing immunoglobulin genes in bats have been limited to a few species within the less-speciose suborder Yinpterochiroptera (e.g. *Pteropus* spp.^12,13^, *Rousettus aegyptiacus*^14^, *Rhinolophus* spp.^15^), leaving the vast majority of species unstudied.

We conducted comprehensive germline genetic annotation and evolutionary analysis of Ig loci in a sampling of vespertilionid bats, and single-cell transcriptome studies of B cells and plasma cells of the big brown bat (*Eptesicus fuscus*), a common North American species and target of infectious disease research, specifically rabies virus for which it is a main reservoir species^16–18^. We report that vespertilionid bats have dual, functional, and distinct IGH loci located on separate chromosomes in their genomes. Our findings identify a novel organization of immunoglobulin genes in mammals and terrestrial vertebrates, and provide initial functional characterization of the impact of these loci on the immune system biology of a pivotal zoonotic disease reservoir.

### Two complete, functional heavy chain genomic loci in *E. fuscus*

We identified two immunoglobulin heavy chain loci in a recent high-quality, long-read genome assembly of *E. fuscus* (NCBI genome assembly DD_ASM_mEF_20220401). Rigorous determination of locus organization was enabled by long-read DNA sequencing permitting full assembly of the complex, repetitive loci in their correct chromosomal localization. The smaller locus (IGH locus A, A-IGH) spans 272 kilobases on chromosome 5; the other (IGH locus B, B-IGH) spans 918 kilobases on the telomeric end of chromosome 24. We annotated the immunoglobulin germline genes of these two parallel immunoglobulin heavy chain loci (**Fig. 1**). The overall structure and orientation of both loci are similar to humans with sequential arrays of variable (V_H_), diversity (D), and joining (J_H_) gene segments followed by multi-exon constant region genes (C_H_). No flanking genes appear to have been transposed as A-IGH is flanked by TMEM212 and CRIP1, both genes known to flank the IGH locus in other mammals, while B-IGH is flanked by MXL.

**Figure 1.**
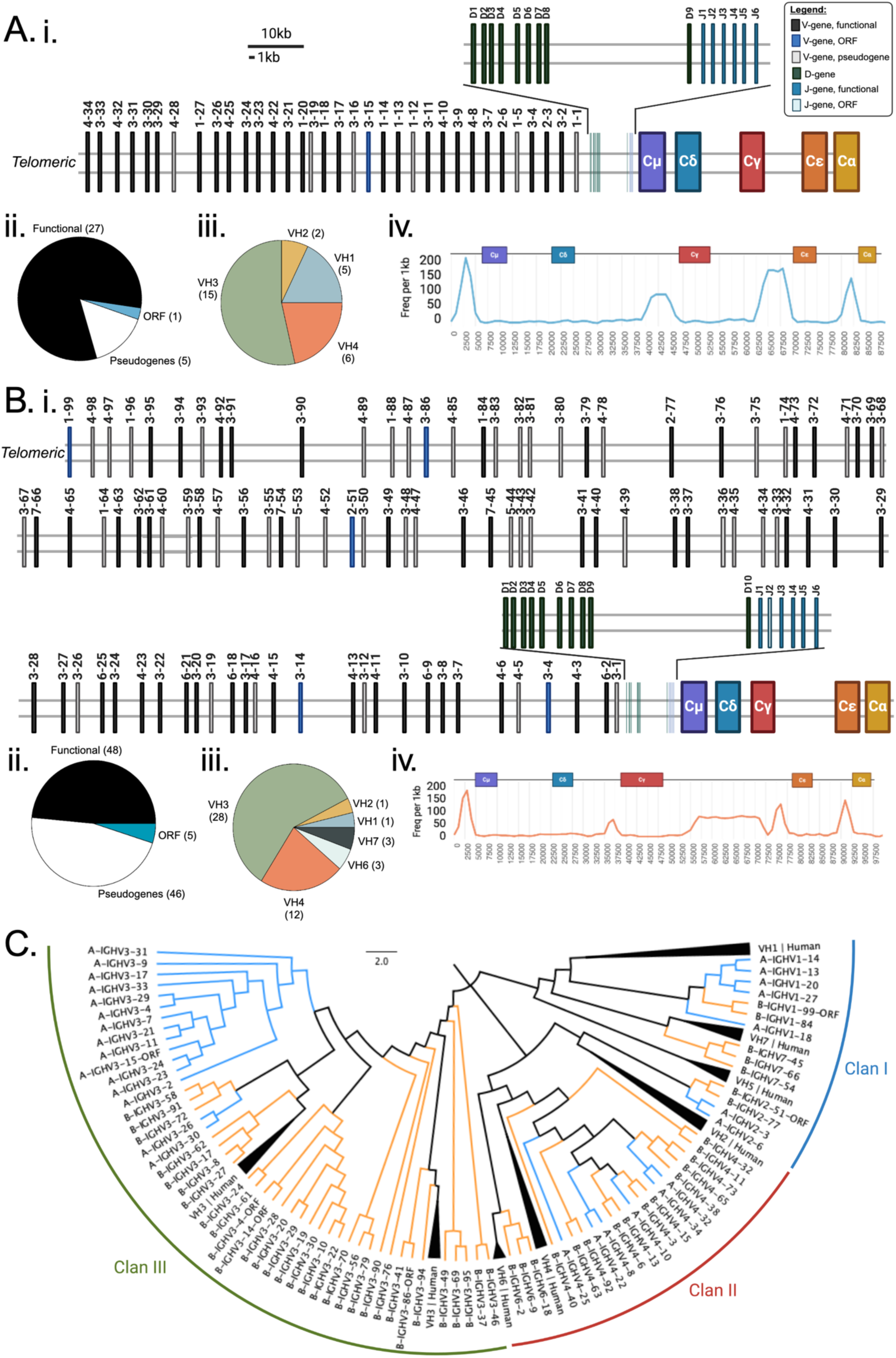
Genomic annotations and analysis for both IGH loci. Genomic annotations and analysis for A-IGH locus contained on contig NC_072477.1 (A) and B-IGH contained on contig NC_072496.1 (B). For each locus, we present (i) a schematic of IG gene organization with an inset for diversity (D) and joining (J) gene, pie charts of (ii) functionality and (iii) gene family distribution for functional variable (V) genes, and (iv) graph of AID hotspot densities per kilobase in the constant gene encoding regions. Nearest neighbor joining tree of functional and ORF V genes for A-IGH and B-IGH with human V genes for reference (C). Created with BioRender.com

Each locus has a single, functional copy of each of the five major mammalian constant region isotype genes, IGHM, IGHD, IGHG, IGHE, and IGHA (**Fig. 1A-B; Supplementary Table 1 - 2**). The order of the IGHCs is consistent with other mammalian species, with IGHM immediately downstream of the IGHJ gene cluster followed by IGHD, IGHG, IGHE, and IGHA. Preliminary annotations suggest that the exonal structure of each gene is conserved within all genes except for IgD. Consistent with previous work^19^, the CH2 exon from IGHD is absent from both gene copies and the hinge H2 exon is fused to the CH3 exon similar to IGHD structures observed in rodents^20^. Additionally, we identified a transmembrane domain previously thought missing in the IGHD gene annotation.

Recombination and isotype switching is initiated by cytidine deamination mediated by the activation induced cytidine deaminase enzyme (AID) which preferentially uses the 5’-AGCT-3’ motif as a hotspot^20–22^. Increased density of canonical mammalian switch motifs immediately upstream of the CH1 domains for each IGHM, IGHG, IGHE, and IGHA in both loci (**Fig. 1A-B**) supports functionality and conserved mechanisms for AID-dependent class switching as observed in other mammalian species. Similar to humans, there is no switch region observed between genes coding for IgM and IgD, consistent with alternative splicing for coexpression of these isotypes^23–25^. Interestingly, B-IGH has a 17.5 kb region with increased switch motif density between B-IGHG and B-IGHE that is not observed in A-IGH which might affect the rate and efficiency of class-switching at this locus.

The IGHJ gene cluster is composed of 6 functional genes on A-IGH and 5 functional genes and one ORF on B-IGH (**Supplementary Table 3**). The canonical di-glycine bulge WGXG motif is encoded in all IGHJ segments and a 23-base pair spacer is observed in the recombination signal sequence (RSS) upstream of each J_H_. All inter-chromosome homologs, except the ORF, share at least 90% nucleotide identity (**Extended Data Fig. 1A**).

Diversity genes were identified by homology as well as RSS searching in the intergenomic space between the most 3’ IGHV and the most 5’ IGHJ. We identified 10 IGHDs on A-IGH and 9 on B-IGH (**Supplementary Table 4**). The IGHDs are more divergent than IGHJs between the loci with only two gene pairs sharing >90% homology (**Extended Data Fig. 1B**). Thus with the duplication, the effective germline IGHD diversity is only slightly smaller than that of the human locus which has 27 germline IGHDs. Notably, the germline IGHDs in A-IGH are longer (mean_A_=18 nt, mode_A_= 19 nt) than those in B-IGH (mean_B_ = 16.67 nt, mode_B_ = 10 nt), potentially contributing to the longer complementarity determining region 3 heavy chain (CDR-H3) loops of sequences originating from this locus, as discussed further below.

While A-IGH encodes just 33 IGHV genes (27 functional, 5 pseudogenes, and a single ORF) (**Supplementary Table 5; Fig. 1A**), the larger B-IGH locus encodes 99 germline IGHVs, of which 48 are functional, 46 are pseudogenes, and 5 are ORFs (**Supplementary Table 6; Fig. 1B**). Interestingly, the percent of pseudogenes is much lower in A-IGH (15%) compared to B-IGH (46.5%) (**Fig. 1A-B**). Both loci contain functional genes from the three major human IGHV clans but differ in the proportions of genes from each of the seven human V_H_ families (**Fig. 1A-C**). As observed with the IGHJ genes, there are highly similar IGHVs between the loci including three genes that have over 90% nucleotide identity (**Extended Data Fig. 1C**). Within clan III (containing the IGHV3 gene family), we observe expansion of functional genes in B-IGH compared to A-IGH. Proportionally, A-IGH is enriched for functional IGHV1 family genes (**Fig. 1A**) while B-IGH contains the only functional IGHV7 and IGHV6 family genes (**Fig. 1B**). These patterns between the loci are consistent with a model in which a full ancestral IGH locus duplicated, followed by separate gene retention, losses and duplications at each locus.

### IGH duplication is common in vespertilionid bats

Comparison of the *E. fuscus* IGHM regions with other bat species revealed that this IGH locus duplication is present across many vespertilionid species (though not all, e.g., *Antrozous pallidus*, data not shown). Each IGH locus forms a separate, well supported clade. Phylogenetic relationships within a given IGHM locus are consistent with the phylogenetic relationships between the species themselves, supporting a single duplication of the IGH locus in the vespertilionid ancestor or very early in the family’s history (**Fig. 2**). All IGHM genes (except *Myotis brandtii* NW005353568, which had too much missing data) have CH1, CH2, CH3, CH4-S and M domains without stop codons. Additionally, all cysteine residues present in the human IGHM are also present in the bats, suggesting the same disulfide bonds that stabilize the structure of the immunoglobulin in humans are present in bats.

**Figure 2.**
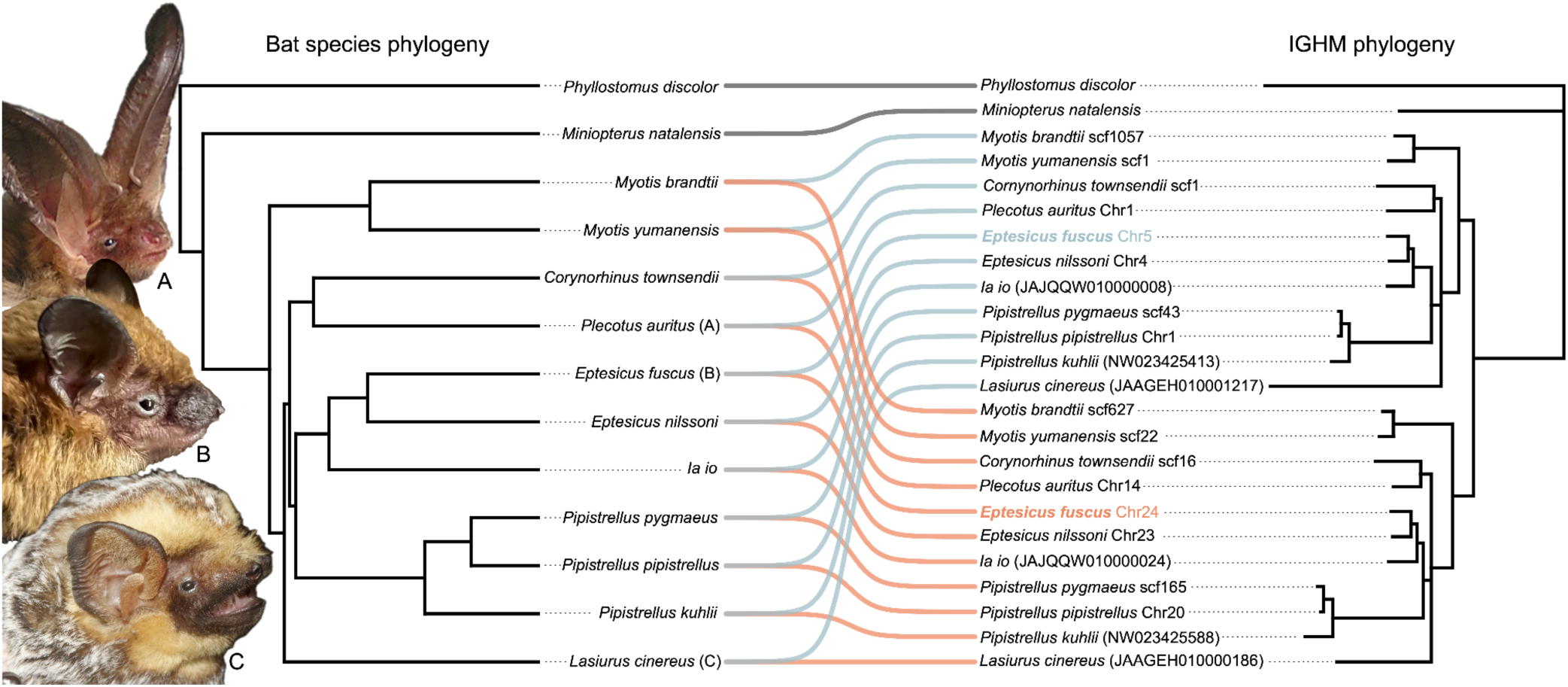
IGHM duplication phylogeny mirrors species phylogeny. The phylogenetic hypothesis on the left is the inferred relationship between the species^60^. The phylogenetic hypothesis to the right is the inferred relationships between IGHM loci. Identifiers of the IGHM phylogenetic hypothesis refer to the chromosome or scaffold on which each IGHM is found. Lines connect the species to its IGHM sequences with blue lines going to A-IGH-like loci and orange lines going to B-IGH-like loci. Bat profiles are A. *Plecotus auritus*; B. *Eptesicus fuscus*; C. *Lasiurus cinereus*. Picture credits are in the acknowledgments.

### Conserved and divergent constant region features related to Fc receptor binding

The key protein features of Ig heavy chain constant regions responsible for contacting Fc receptors to initiate effector functions are well described^26,27^. To understand potential for locus specific effector functions of IGHG, we looked for conservation of features that mediate Fc receptor binding: (1) the conserved N-glycosylation site at Asn^297^; (2) CH2 region residues near the lower hinge found in IgG1, IgG3 and IgG4 (Leu^234^ Leu^235^ Gly^236^ Gly^237^); (3) CH2 BC loop Asp^265^; and (4) CH2 FG loop (Ala^327^ Leu^328^ Pro^329^ Ala^330^ Pro^331^). The IGHGs from both *E. fuscus* loci contain the conserved N-glycosylation site, conserved Asp^265^, and the canonical CH2 FG loop, only with an Iso^328^ instead of a Leu^328^. The cytosolic tails are also highly similar to humans containing the canonical motifs (DYXNM and SSVV^23^ [SSVA in *E. fuscus*]) for intracellular signaling.

The hinge length and sequence play an important role in the function of different IgG sub isotypes in humans. We found a single hinge exon (H) for each IGHG similar to IgG1, IgG2, and IgG4 in humans, but the sequence differs between A-IGHG and B-IGHG and between *E. fuscus* and humans. Both bat IGHG H exons encode only two cysteines similar to IGHG4 H in humans, however the lengths of the bat IGHG H regions are more similar to IGHG1 H and IGHG3 H2-H4 genes. A-IGHG H possesses the run of TPPPC found in the IGHG3 H2-4 genes, while B-IGHG H has TTTPC which is more similar to the TTHTC of IGHG3 H1 gene. Additionally, in the human CH2/ lower hinge region, the flexibility of residues Pro^232^–Pro^238^ is critical for Fc receptor binding^26^; both A-IGHG and B-IGHG encode a Cys^232^ instead of a Pro^232^ and a Thr^234^ instead of a Leu^234^, suggesting a modified interaction compared to human constant region binding to Fc receptors.

### *E. fuscus* has retained lambda and lost kappa light chains

Antibodies are made up of heavy and light chains. In mammals, light chains are of two types, kappa (κ) or lambda (λ). Salmon and rainbow trout, both species of fish that have a duplicated IGH locus, also express multiple types of light chains^28,29^. In contrast, birds have a single light chain, lambda (λ)^30–32^ which is postulated to be the result of genome size restraints linked to their high metabolic rate and adaptation to flight. In light of the IGH duplication, we asked whether *E. fuscus* has two light chain loci, as do other mammals and fish, or a single light chain locus, similar to birds. We identified a single light chain lambda locus (**Fig. 3A; Supplementary Table 7-9**). Neither the NCBI eukaryotic genome annotation pipeline^33^, IgDetective^34^ nor initial BLAST^35^ searching detected any kappa variable, joining, or constant-like genes in the current assembly. In humans, the IGK locus is on chromosome 2 near the centromere, flanked on the telomeric side by RPIA in reverse orientation and EIF2AK3 in forward orientation. In *E. fuscus*, these genes are found in close proximity on chromosome 9. A focused search for IGK-like genes from 3.5Mb upstream of RPIA to 1Mb downstream of EIF2AK3 revealed three IGKJ but only one was preceded by two overlapping RSS sequences, one with a 12bp spacer and the other with a 23bp spacer (**Fig. 3B**). However, no IGKC was found in between the IGKJ and RPIA genes, nor were any IGKV genes found in the 5Mb region, suggesting loss of the IGK locus in the evolutionary history of *E. fuscus*. We additionally searched single cell data presented below for evidence of a second light chain but found only IgL CDR3 in 89.8% of B lymphocytes and 96.7% of the cells for which we have complete, productive IGH, with no evidence of kappa light chains. Together, these data support that *E. fuscus* has no functional or expressed kappa light chains, consistent with previous work^36^.

**Figure 3.**
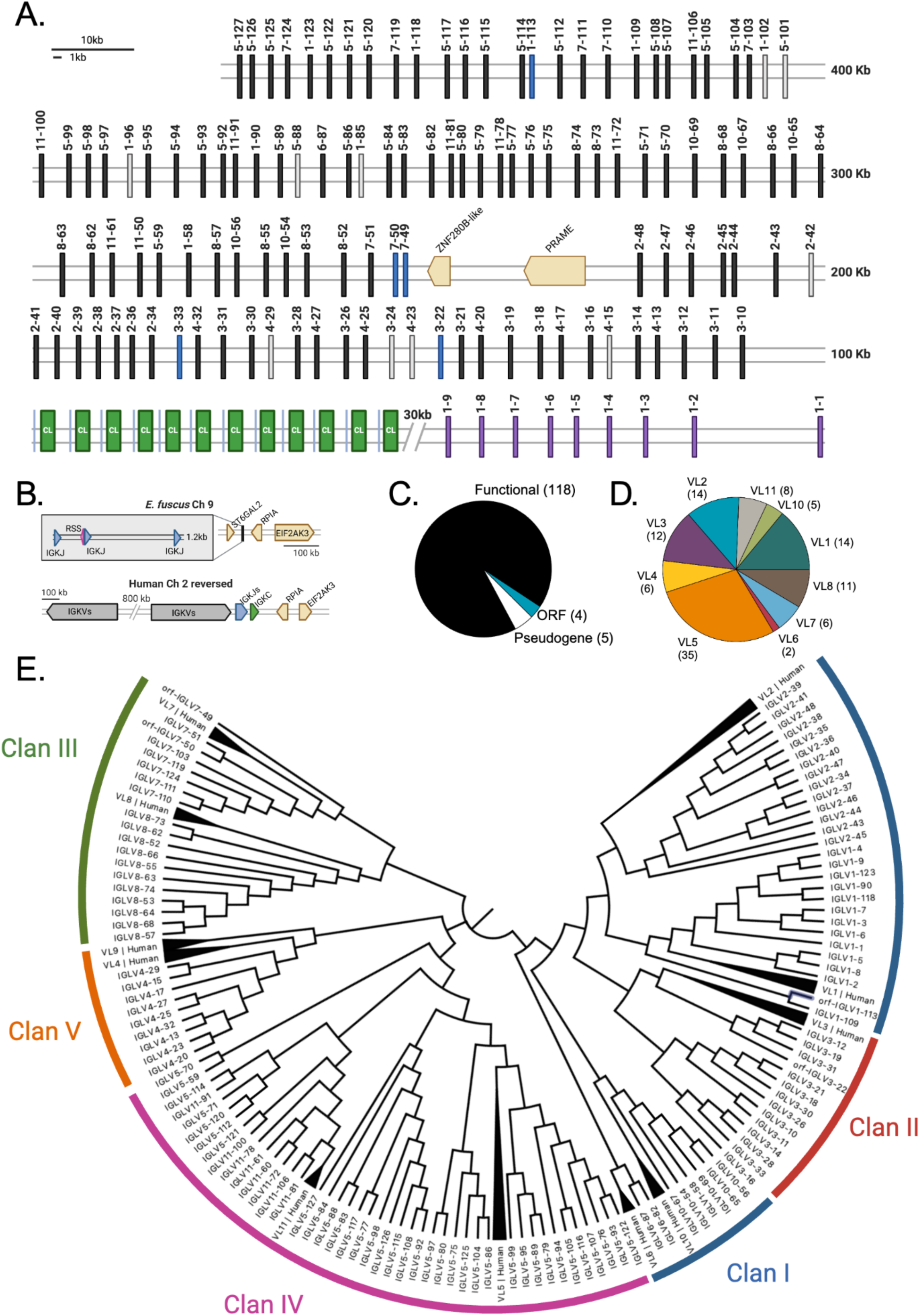
Annotation of single lambda-like light chain locus. (A) Schematic of IG gene organization for lambda locus contained on contig NC_072495.1 (B) Schematic of vestigial IG kappa locus on chromosome 9 (contig NC_072481.1) with the human locus for comparison. Pie chart of (C) functionality and (D) family distribution for lambda variable genes. (E) Nearest neighbor joining tree of functional and open reading frame (ORF) V genes with human V genes for reference and human clans. Created with BioRender.com

The *E. fuscus* lambda locus on chromosome 23 (contig NC_072495.1) spans only 506 kilobases despite encoding 127 variable (Vλ, Table 7) genes, 12 joining (Jλ, Table 8), and 12 constant (Cλ, Table 9) genes. The locus is compact compared to the human kappa locus that encodes 116 genes but spans 1.9 megabases (Mb). The organization of Jλ and Cλ genes mirrors other species, such that each Cλ gene is preceded by one Jλ gene, forming 12 J-C gene clusters (**Fig. 3A**). Each Jλ gene is flanked on the 5’ end by an RSS with a 12-base spacer and the Vλ genes with a 3’ RSS with a 23-base spacer as is observed in other species. Notably, the locus encodes nine functional Vλ genes downstream of the Jλ - Cλ gene cassettes in an inverted orientation (**Fig. 3A**). This organization is also seen in the equine lambda locus^37^.

We found Vλ genes from all five human clans and all V families except Vλ9 in *E. fuscus* (**Fig. 3C, 3E; Supplementary Table 7**). Of the 127 Vλ genes, 118 are functional, 4 are open reading frames, and 5 are pseudogenes (**Fig. 3D**), nearly double the number of functional genes found in the human lambda and kappa light chain loci combined. In analyzing the rearranged repertoire, all functional Vλ genes, three ORFs, and a pseudogene, IGLV1-102^P^, were identified in productive rearrangements. Sequences using the inverted V genes comprised 6% of the total repertoire, indicating that functional rearrangement does occur despite the non-canonical orientation.

### Distinct B cell phenotypes within *E. fuscus* spleen

Different B cells subsets serve different functions during an immune response, as do the antibodies they produce. To investigate *E. fuscus* B cell and plasma cell phenotypes, we performed single cell RNA-seq on cryopreserved splenocytes from four individual bats using the 10x platform. After pre-processing, batch correction, and removal of low quality cells, 62,747 cells were analyzed. Using canonical transcriptional markers present in the current genome annotation, we identified clusters corresponding major immune cell types (**Fig. 4A**). We identified two B lymphocyte lineage clusters, one enriched for B cell markers (i.e., *MS4A1* and *CD19*) associated with naive and memory B cells while the other was enriched for plasma cell markers (i.e., *XBP1, MZB1, PRDM1, SDC1 and JCHAIN*) (**Fig. 4B**).

**Figure 4.**
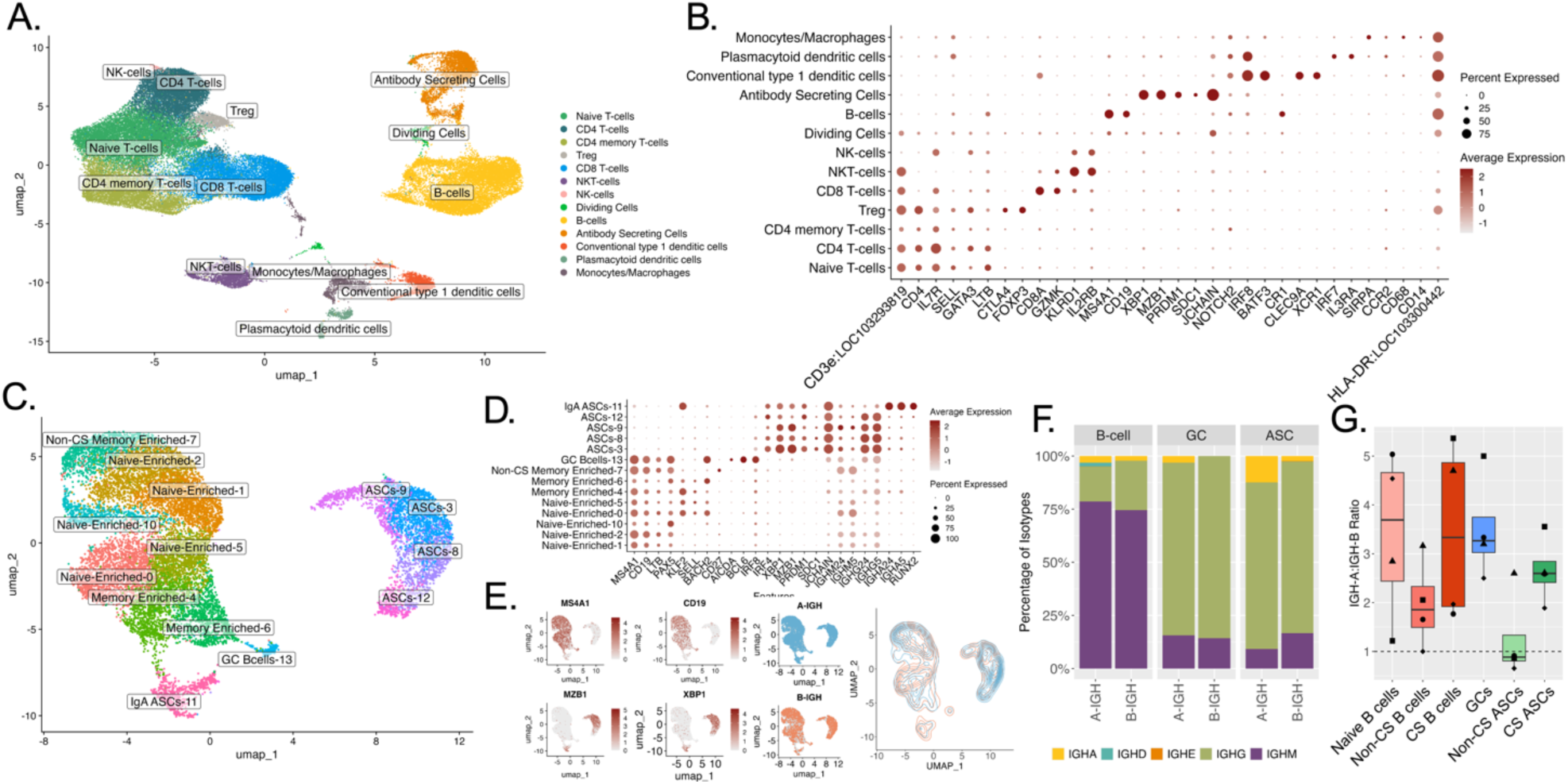
Splenic single cell transcriptomics reveal distinct B cell phenotypes. (A) UMAP representation of 62,747 single cell transcriptomes from four *E. fuscus* bats’ (*n* = 4) spleens. Cell clusters were annotated and grouped according to major cell type classification based on canonical marker genes. (B) Dot plot of the main marker genes used to define each cell type. Dot size is proportional to the percentage of cells with detectable expression of the indicated gene and dot color is indicative of the average expression value for the indicated gene, scaled across all identified clusters. (C) UMAP representation of subclusters of 14,618 cells within the B lymphocyte compartment after doublet exclusion. (D) Dot plot of the canonical B lymphocyte lineage and activation marker genes used for phenotype classification. (E) Feature plot of log-normalized expression of marker genes used to distinguish B cells from antibody secreting cell phenotype cells. UMAP highlighting 5,010 single cells with full length IGH B cell receptor (BCR) assemblies from A-IGH (blue) and B-IGH (orange). Density plot of cell with A-IGH (blue lines) and B-IGH(orange lines). (F) Stacked bar plot of the percent of each isotype of BCRs from A-IGH and B-IGH for B cells, germinal center (GC), and antibody secreting cells (ASCs). (G) Box plot of ratio of A-IGH:B-IGH for each cell phenotype. Line represents the mean and ratio for each individual (*n* = 4) plotted with different shapes. Dashed line at ratio of 1. Created with BioRender.com

In humans, there are well-defined subsets of B cells found in the spleen with different activation states as well as stages of development^38^. Many subsets described in humans have relatively similar counterparts in mice^39^, but are less well understood in other mammals. Previous studies have identified B lymphocytes in scRNAseq analysis in various bat species^40–42^ but functional B cell subsets have not been fully defined, in part due to a lack of Ig gene annotations. Therefore, we undertook a focused analysis of B lymphocyte lineage subsets in *E. fuscus*. We identified a cluster analogous to germinal center (GC) cells that express canonical human markers such as *BCL6, BACH2* and *AICDA* (**Fig. 4D**). Naive, transitional, and memory B cells showed less distinct cluster separation. This is consistent with other single cell studies in various bat species^40–42^ in which clusters were mixed or defined by non-canonical genes of unknown cell phenotype specificity. We found clusters enriched for memory-associated genes (e.g., *NOTCH2*^43^*, VIM, ZBTB32*^44^*, CD27*), and others enriched (i.e., ≥ 50%) for B cells with low somatic hypermutation (SHM) frequencies (< 1.5%), and expression of IGHM/IGHD isotypes presumed to be naive B cells (**Extended Data Fig. S2**).

Further genome annotation curation and experimental validation will be needed to fully define the gene expression patterns of naive and memory B cell populations, but we adopted a functional classification of naive B cells as those expressing IGHM or IGHD without mutation (SHM <1.5%). We grouped the remaining somatically mutated cells into non-class switched B cells (non-csB-cells) or class switched B cells (csB-cells) based on their expression of IGHM/D or other isotypes, respectively.

In contrast, plasma cell phenotype clusters were relatively homogeneous with elevated expression of canonical markers (i.e., *IRF4, MZB1, JCHAIN, XBP1*) except for a cluster of IgA class-switched plasma cells. This cluster is enriched for *RUNX2*, a transcription factor shown to induce IgA class switching in mature B cells during BCR signaling^45–48^. We divided the plasma cells into class-switched antibody secreting cells (csASC) and non-class switched antibody secreting cells (non-csASC).

### Rearrangement and usage of the dual IGH loci in *E. fuscus*

Having determined that numerous vespertilionid bats have dual heavy chain loci with different sizes and gene segment composition, we sought to answer key questions about the use of these loci in B cells and plasma cells, specifically: 1) whether there is allelic exclusion preventing each B cell from making two different heavy chains; 2) whether inter-locus V-D-J rearrangements occur; 3) how rearranged VDJ products of the two loci differ in gene segment and CDR-H3 characteristics; 4) whether locus usage is correlated with the B cell or plasma cell phenotype; 5) whether class switching or SHM processes differ between the two loci; and 6) whether there is evidence of selection favoring particular IGHV genes in antigen-experienced B cells and plasma cells.

### Strong allelic exclusion of the dual IGH loci and evidence for A-IGH rearranging before B-IGH

Allelic exclusion is the process that ensures each B cell expresses only one heavy and one light chain sequence despite having two chromosomal copies of each locus, and two different light chain loci in most mammals. It is considered to be an important mechanism for controlling autoreactivity that could result if B cells expressed multiple antigen receptors specifically self-reactive and pathogen-specific receptors on the same cell which would allow for maturation to occur resulting in a cell where response to a pathogen could trigger autoimmunity. We hypothesized that allelic exclusion in bats with dual IGH loci would ensure that only a single IGH locus would be functionally rearranged to give an in-frame expressed protein product in each B cell or plasma cell. Using custom primers, we generated enriched and sequenced BCR libraries for single cells and assembled IGH rearrangements with CDR-H3s for 9,922 (68%) of the 14,618 B lymphocytes identified in the gene expression dataset. After removing low confidence assemblies, we found only 198 cells appearing to have two expressed in-frame CDR-H3 transcripts, or 2.0%. Given potential artifacts of single cell sequencing such as incomplete removal of doublet events or RNA from lysed cells associating with the surface of other B cells, we interpret these findings as most consistent with allelic exclusion acting at the dual IGH loci.

To address IGH locus usage in B cells of different cellular phenotype, and to assess whether one locus is rearranged before the other in the formation of the naive B cell repertoire, we assembled full length BCR heavy sequences from 5,010 single B cells or plasma cells. Both A-IGH and B-IGH are used with productive rearrangements in all phenotypic cell subsets (**Fig. 4E**). Clusters 0, 1, 2, 5, and 10 contain predominantly naive B cells expressing IgM, while IgG dominates the GC and ASC clusters (**Fig. 4F**; **Extended Data Fig. 2**). We assume that B cell precursor IGH V-D-J rearrangement attempts are successful in forming an in-frame product one-third of the time because the V and J genes must be in the same reading frame in the final rearrangement, and exonuclease chewback of V, D and J segment ends together with random base addition by the terminal deoxynucleotidyl transferase at the V-D and D-J junctions can alter the J reading frame relative to the V reading frame. Therefore, sequential attempts to rearrange both copies of A-IGH and then B-IGH would result in a ratio of A-IGH: B-IGH usage in naive cells of approximately 2.25, whereas random selection of loci to rearrange would result in a 1:1 ratio. The mean ratio of A-IGH to B-IGH locus usage we observed in naive cells is 3.41 (**Fig. 4G**). This bias supports a model of ordered rearrangement and allelic exclusion such that A-IGH rearrangements are attempted first, and only if no productive V-D-J rearrangement is obtained is rearrangement attempted at B-IGH, resulting in a larger percentage of naive cells with A-IGH BCRs compared to B-IGH. Other possible explanations for this ratio of A-IGH to B-IGH could include strong negative selection against B cells expressing the B-IGH locus in the developing B cell.

### No evidence of cross loci rearrangement

We assessed whether VDJ rearrangement occurs between loci by determining the chromosome of origin for each BCR by looking for agreement between the origin of the V gene and the J gene. The origin of the C-gene was not used due to lack of polymorphisms upstream of the enrichment primers retained after library preparation. The D gene origin was also not considered due to the greater uncertainties in reliably identifying sequences derived from D genes in rearrangements, and the level of homology across chromosomes for certain D-genes. Over 95% of sequences had a V gene and a J-gene from the same locus. We found similar patterns and no evidence of cross-loci rearrangement in the sequenced bulk BCR repertoires (*data not shown*).

Although we cannot fully rule out the possibility of inter-locus recombination, it is likely that the apparent locus mismatches are due to errors in germline gene calling related to the high homology between certain V_H_, D, and J_H_ genes between the loci (**Extended Data Fig. 1**), and our limited knowledge of allelic variation between *E. fuscus* individual animals. We therefore used the V gene locus alignment to define the origin of each IGH sequence, as the V region is longer than the J region, and therefore far more likely to have the correct locus call.

### Differences in VDJ rearrangements from the dual IGH loci

As discussed above, the germline repertoire is largely similar across different loci but exhibits notable differences that likely contribute to the diversity of expressed repertoires. We hypothesized that rearranged VDJs from different IGH loci exhibit distinct V gene segment usage and complementarity-determining region 3 (CDR-H3) features. To capture a representative sample, expressed immunoglobulin rearrangements were enriched for IgM and IgG isotypes using 5’-RACE cDNA from flash frozen splenic tissue of six animals yielding a total of 307,312 productive BCR sequences for the following analysis.

Genes from the VH3 family are the most common in both the IgM and IgG repertoires for both loci (**Extended Data Fig. 3A-B**). However, for A-IGH, VH3 family gene usage significantly decreases in IgG compared to IgM (p < 0.05 by clone-wise non-paired Wilcoxon rank-sum test), while VH1 family gene usage significantly increases (p < 0.05 by clone-wise non-paired Wilcoxon rank-sum test). For B-IGH, there is almost no VH1 usage and heavy usage of VH7 family genes which significantly increases in IgG clones compared to IgM (p < 0.05 by clone-wise non-paired Wilcoxon rank-sum test). These data demonstrate differences in the V gene usage during generation of the naive repertoire. Furthermore, differences in IGHG gene usage support locus specific antigen-driven selection.

Junctional diversity was assessed by comparing the length V-D (N1) and D-J (N2) regions between chromosomes for IGHM associated sequences. The N1 region is significantly shorter for A-IGH (mean_A-IGH_ = 3.1 nucleotides) compared to B-IGH (mean_B-IGH_ = 3.4 nucleotides; p < 0.05 by clone-wise non-paired Wilcoxon rank-sum test) while N2 region is not significantly different (mean_A-IGH_ = 1.76 nucleotides; mean_B-IGH_ = 1.78 nucleotides; p = 0.5 by clone-wise non-paired Wilcoxon rank-sum test; **Extended Data Fig. 3C**). Overall, *E. fuscus* shows fewer N nucleotide additions compared to humans that average 7.7 and 6.5 nucleotides N1 and N2 respectively^49^. These limited nucleotide additions likely function to complete partially encoded codons rather than introducing new non-templated amino acids as observed in humans. Thus, the junctional diversity of these bats appears to be constrained compared to humans^49^ leading to a more germline-focused IGHM repertoire.

The length of CDR-H3s from A-IGH is significantly longer than B-IGH for both IGHM (mean_A-IGH_ = 10.2 amino acids, mean_B-IGH_ =9.3 amino acids not including the initial cysteine and terminal tryptophan residues; p <0.01 by clone-wise non-paired Wilcoxon rank-sum test) and IGHG (mean_A-IGH_ = 10.6 amino acids, mean_B-IGH_ =9.1 amino acids not including the initial cysteine and terminal tryptophan residues; p <0.01 by clone-wise non-paired Wilcoxon rank-sum test; **Extended Data Fig. 3D**), owing primarily to the longer IGHDs encoded by the A-IGH locus.

### Evidence of selection pressure on IGH locus usage in B cell subsets

To assess for selection pressures affecting A-IGH or B-IGH usage in different B cell subsets, we compared the ratio of locus usage across the major B cell and plasma cell phenotypic groups, compared to naive B cells. The A-IGH:B-IGH ratio in non-csB cells is lower compared to naive B cells and it remains relatively unchanged in csB cells (**Fig. 4G)**. The A-IGH bias is only lost in non-csASC, mean = 1.26, which is significantly lower than in naive cells (p = 0.05 by non-paired Wilcoxon rank; **Fig. 4G)**. A Chi-square test comparing the frequency of A-IGH and B-IGH usage across the different B cell subsets showed a highly significant difference (p < 0.001), indicating a strong association between B cell phenotype and locus usage.

### Decreased class switching in the B-IGH locus

Notably, IgA csB cells and csASCs disproportionately used the A-IGH locus, indicating that this locus may have particular importance for mucosal humoral responses dependent on IgA (**Fig. 4F)**. Further, ASCs using B-IGH were more likely to be non-class switched, expressing IgM, compared to those using A-IGH (**Fig. 4F**). Decreased class switching to IgA in the B-IGH locus could be related to the unusual, oversized switch region between IGHG and IGHE compared to A-IGH (**Fig. 1B**), but we observed relatively few IgE-expressing B cells or ASCs. Therefore, despite the greater diversity of IGHV genes in the B-IGH locus, it is more often used in non-csASCs such as those that are described in other mammals as being part of primary and extrafollicular humoral responses^50–52^. Overall, these data support a model in which the B-IGH locus is less efficient at class-switching, potentially due to decreased recruitment or accessibility of AID, or that B cells using the B-IGH locus are less likely to enter immunological niches promoting class switching such as the germinal center.

### Decreased somatic hypermutation in the B-IGH locus

To further evaluate functional differences between the two IGH loci, we measured the frequency of SHM changes in VDJ rearrangements. In mammals, SHM is due to focused recruitment of the AID enzyme to the IG loci, followed by DNA repair processes triggered by initial cytidine deamination, and is the basis for affinity maturation in which B cells expressing higher affinity for antigen preferentially expand within the stimulated clone. To look for potential SHM differences between loci, we analyzed IGH sequences with complete VDJ assemblies (5,010 of the single B lymphocytes, 34% of total B cells identified, 50% of those with a CDR3). Rates of somatic hypermutation ranged up to 22.7%. As expected, we observed the lowest mean SHM in IgM sequences (1.8%) and higher mean rates in IgA and IgG (4.8% and 4.1%, respectively) in the B cell clusters. The SHM rates were not significantly different in ASCs compared to non-naive B cells for all isotypes except IgM (p < 0.01, clone-wise non-paired Wilcoxon rank-sum test).

For A-IGH, class-switched cells had significantly higher levels of SHM compared to non-class-switched cells within ASCs (p<0.001, clone-wise Kruskal-Wallis rank sum test followed by Dunn’s multiple comparisons) (**Fig. 5A, left panel**). This is consistent with what is observed in humans and suggests similar mechanisms likely result in class switching early in activation resulting in low mutation IgM. However, this is not observed when comparing cs to non-cs B cells for A-IGH (p=0.50 by clone-wise Kruskal-Wallis rank sum test followed by Dunn’s multiple comparisons). For B-IGH, however, the SHM of non-class switched cells is higher than class switched B-cells and there is no difference in ASCs (**Fig. 5A, right panel)**.

**Figure 5.**
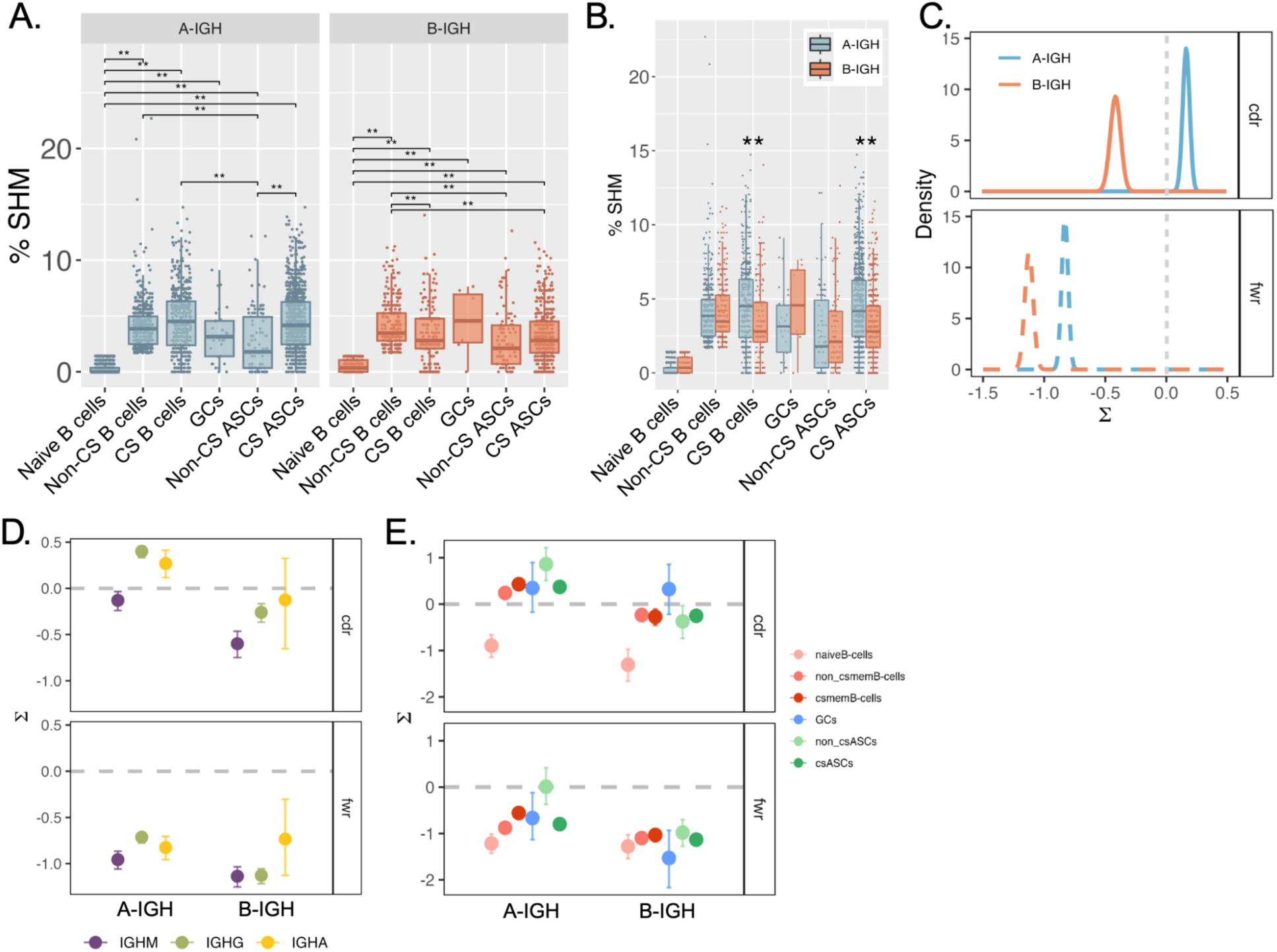
Levels of SHM vary between and within IGH loci. (A) Distribution of percent nucleotide SHM of IGHV segment for each clone grouped by phenotype and locus. (B) Distribution of percent nucleotide SHM of IGHV segment for A-IGH (blue) and B-IGH (orange) clones grouped by cell phenotype. (C) Density plot of selection scores (sigma, Σ) of the CDRs (solid line, top panel) and FWR (dashed line, bottom panel) for IGHV segments from A-IGH (blue) and B-IGH (orange) clones. Threshold for positive selection gray dashed line (Σ = 0). (D) Plot of mean with confidence interval for the selection scores of CDR (top panel) and FWR (bottom panel) grouped by locus for IgM (purple), IgG (green), and IgA (yellow). (E) Plot of mean with confidence interval for selection scores of CDR (top panel) and FWR (bottom panel) grouped by locus for each cell phenotype. Comparisons between groups were performed with the Wilcoxon rank-sum test. **p < 0.05. Graphs generated in R and figures created with BioRender.com

This supports the hypothesis that B-IGH cells remain non-class switched through multiple rounds of SHM compared to class switched cells. Alternatively, once class switching has occurred, the recruitment of AID is impaired. There is no significant difference in SHM rate of csB-cells compared to csASCs within either locus. The level of SHM in non-csB-cells is significantly higher than in non-csASCs for both B-IGH and A-IGH (p < 0.001 by clone-wise Kruskal-Wallis rank sum test followed by Dunn’s multiple comparisons) conforming to canonical trends in SHM.

The SHM rates in csB-cells are significantly higher in A-IGH sequences compared to B-IGH (p < 0.001 by clone-wise non-paired Wilcoxon rank-sum test) as well as in csASCs (p < 0.001 by clone-wise non-paired Wilcoxon rank-sum test), but there was no significant difference in non-class switched cells or GCs (**Fig. 4B**). These findings and statistical significances were confirmed by permutation tests (**Extended Data Fig. 4**). Together, these results support the hypothesis that the ability of AID to access each locus is different.

### Greater evidence for selection pressure on SHM changes in A-IGH

Antigen experience shapes the B cell receptor repertoire through selective expansion of B cells with high-affinity receptors. Positive selection for improved antigen binding occurs mainly in the complementary-determining regions (CDRs) of an antibody, while negative selection against destabilizing mutations acts most strongly on the intervening structural framework regions (FWRs). Selection scores (sigma, Σ) are higher in complementary determining region (CDR) compared to the framework regions for both A-IGH and B-IGH (**Fig. 4C**). This supports similar mechanisms of affinity maturation in both loci of *E. fuscus* as observed in other species in which the antigen contacting regions are under positive selective pressure while other regions are not. However, positive selection (Σ > 0) only reached significance for the CDRs of A-IGH, but not B-IGH (**Fig. 4C**). This is likely due to the overall lower levels of SHM in B-IGH sequences resulting in the inability to detect positive selection above background without the aid of a specialized model to detect mutational hot and cold zones as is used in humans. Furthermore, this evidence of positive selection is only found in IgG and IgA, but not IgM for A-IGH sequences (**Fig. 4D**). Lastly, the only B-IGH sequences that demonstrate positive selection are associated with GC B cells. As stated above, this is likely due to the overall lower levels of somatic hypermutation in B-IGH sequences, except GC B cells (**Fig. 4B**). One possible explanation for these data would be if A-IGH more easily engages AID compared to B-IGH. The data could also suggest a propensity for B cells using A-IGH to engage in follicular responses while B cells using B-IGH may participate more often in extrafollicular responses.

### A novel strategy for mammalian humoral immunity

Our analysis of high-quality long-read chromosome level assemblies for *E. fuscus* and other vespertilionid bats identified novel IGH locus duplications, highlighting the limited extent to which these complex, repetitive loci have been explored in mammals and other vertebrates. These bats share many common features of humoral immunity with other mammals such as allelic exclusion, somatic hypermutation, evidence of antigen-driven selection, and class switching, but also provide evidence for unique functional specialization of the two IGH loci.

The A-IGH and B-IGH germline loci differ notably in size, gene segment complexity, D gene lengths, and class switch region distributions, but both have similar constant region composition and can produce IgM, IgD, IgG, IgE and IgA. VDJ rearrangements from the shorter, less V gene rich A-IGH locus are more than 3-fold more common than rearrangements from B-IGH in naive B cells, supporting a model in which rearrangement of A-IGH is attempted at both chromosome copies in developing B cell precursors prior to attempting to rearrange the B-IGH locus. A-IGH rearrangements have longer CDR-H3s than those of B-IGH, adding diversity for this key protein loop that is often involved in antigen binding. The greater diversity of B-IGH V segments encoding CDR-H1 and CDR-H2 loops is initially deployed in a smaller fraction of the naive B cell repertoire in these bats.

Both IGH loci are used in antigen-experienced B cells and plasma cells whose BCRs contain SHM changes with or without class-switching, but A-IGH locus rearrangements have consistently higher SHM and are more often class switched. These data could indicate that B cells or plasma cells expressing A-IGH rearrangements are more likely to engage in germinal center reactions in the bats, or could point to an overall deficiency in the ability of the B-IGH locus to recruit the AID enzyme complexes required for both SHM and class-switching. Functional consequences of these divergent loci are likely to be significant; for example, in these bats most IgA, the key antibody isotype for mucosal immunity, derives from the A-IGH locus with its more limited repertoire of IGHV genes. Nonetheless, usage of the B-IGH locus increases in antigen-experienced B cells and plasma cells, suggesting that the greater diversity of IGHV gene segments in this locus provide greater possibilities for recognition of foreign antigens and potential pathogens. We speculate that in these bats the fraction of their secreted antibody populations derived from B-IGH, characterized by less-frequent class-switching and lower SHM, but higher gene segment diversity, could provide a broad, lower-affinity arm to their humoral immune responses, complemented by A-IGH-derived antibody responses that have higher SHM frequencies, greater evidence of antigen-driven selection for increased affinity, and higher rates of class-switching to IgG or IgA for potentially altered effector functions.

The combination of these humoral immunity types could contribute, together with the innate immune antiviral adaptations reported in many bat species^53,54^, to their greater resistance to viral pathogens and their roles as vectors for asymptomatic or less-symptomatic transmission of zoonotic diseases to humans. We note that the duplication and retention of two IGH loci is particularly striking in bats, because flying species typically have genomes of reduced size^55^; consistent with this, bats have smaller genomes than most other mammals^56^. Thus, there may be a significant selective advantage to having two functional IGH loci that outweighs the pressures to reduce genome size.

The finding of two functional IGH loci in vespertilionid bats is the most striking example of emerging themes in bat immunity research: that bats share the same molecular components of immunity with their mammalian relatives but show bat-specific adaptations; and that gene family evolution is critical for bat immune adaptation^57,58^. It also emphasizes the differences between bat species, as we found no evidence of the IGH duplication outside of the Vespertilionidae, the same family in which recent work has found a duplication of an important antiviral protein, protein kinase R^59^.

Further testing of the functional roles of the A-IGH and B-IGH loci will benefit from studies of vaccination and infectious challenges in vespertilionid bats. Analysis of the dual IGH loci should provide a unique internally-controlled opportunity to elucidate the fundamental genetic and epigenetic mechanisms governing the formation and evolution of B cell receptor repertoires in intact primary B cells.

## Acknowledgments

We would like to thank Yana Safonova for insightful discussions regarding the presented research. We are grateful to the following organizations for funding the work: National Science Foundation IOS (2032157; HKF), National Institutes of Health NRSA T32 (T32OD011121, TP), National Institutes of Health NIAID (5 R21 AI169548; HKF), Life Sciences Research Foundation Fellowship (HKF), Open Philanthropy Project (HKF), Stanford Woods Institute for the Environment Environmental Venture Program (SDB; HKF), Stanford Center for Computational, Evolutionary and Human Genomics Postdoctoral Fellowship (HKF), National Institutes of Health grants ‘Molecular and Cellular Immunobiology’ (5 T32 AI07290; HKF), Stanford School of Medicine Dean’s Postdoctoral Fellowship (HKF), funds from the Stanford Pathology Department and an endowment from the Crown foundation (SDB). This work was supported by funding to The Viral Emergence Research Initiative (viralemergence.org) from the U.S. National Science Foundation, including NSF BII 2021909 and NSF BII 2213854 (HKF). We thank the skilled staff of the Stanford Clinical Genomics Laboratory for assistance with sequencing experiments. The authors would also like to thank the invaluable contributions and animal care assistance supported by members of the Poxvirus and Rabies Branch and the Comparative Medicine Branch at CDC, Atlanta, GA.

## Author contributions

Conceptualization: TP, SDB, HKF; Methodology: TP, AR, AM, PK, JAE, SDB, HKF; Software: TP, AM, PK; Formal analysis: TP, AR, HKF; Samples: JAE, CLH; Data Curation: TP, AR, HKF; Writing-Original Draft: TP, SDB, HFK; Writing-Review & Editing: TP, AR, AM, PK, JAE, CLH, SDB, HFK; Visualization: TP, AM; Supervision: SDB, HFK; Funding acquisition: JAE, SDB, HFK.

## Disclaimer

Use of trade names and commercial sources are for identification only and do not imply endorsement by the US Department of Health and Human Services or the Centers for Disease Control and Prevention. The findings and conclusions in this report are those of the authors and do not necessarily represent the views of their institutions.

## Additional information

**Supplementary information is available for this paper.**

**Correspondence and requests for materials should be addressed to** H.K. Frank and Scott Boyd

## Data availability

IG gene annotations can be found in Supplementary Tables 1-9. All raw sequencing reads have been deposited in NCBI’s Short Read Archive (BioProject ####). The GenBank IDs for all genome assemblies used for phylogenetic analysis listed in Supplementary Table 1. Genome assembly and non-IG gene annotations for *E. fuscus* are available through GenBank (https://www.ncbi.nlm.nih.gov/datasets/genome/) under the GenBank ID provided in **Methods Table 1**.

Photo credits (Fig. 2): A. “brown long-eared bat (Plecotus auritus)”

<https://www.flickr.com/photos/wolf_359/123404678/> by wolf_359 is used under a CC BY-NC-SA 2.0 license. B. “Big brown Bat (Eptesicus fuscus)”

<https://www.inaturalist.org/observations/145464659> by misspt is used under a CC BY-NC 4.0 license. C. “North American Hoary Bat (Lasiurus cinereus ssp. cinereus)”

<https://www.inaturalist.org/photos/18602149> by juancruzado is used under a CC BY-SA 4.0 license. We removed the background from all photos; licensors make no endorsements.

---

**Extended Data Figure 1.**
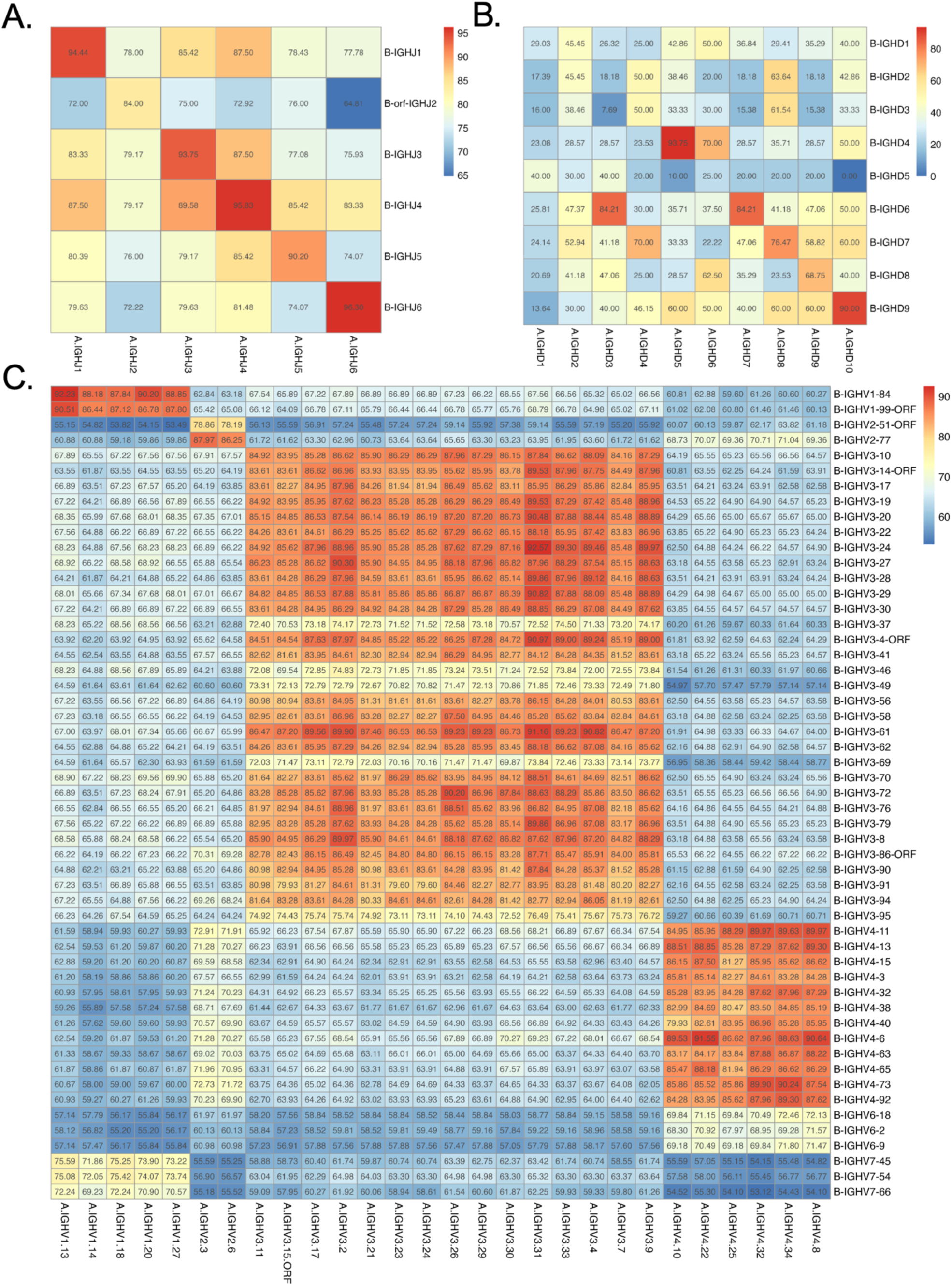
Levels of inter-locus homology varies across germline genes. Heatmaps of the distance matrix of (A) joining genes, (B) diversity, and (C) variable genes from A-IGH compared to B-IGH where the color and number indicate the percent nucleotide identity for the given pair of V genes. MUSCLE alignment with iterations (n = 10) clustered with neighbor joining and CLUSTALW sequence weighting scheme. Created with BioRender.com

**Extended Data Figure 2.**
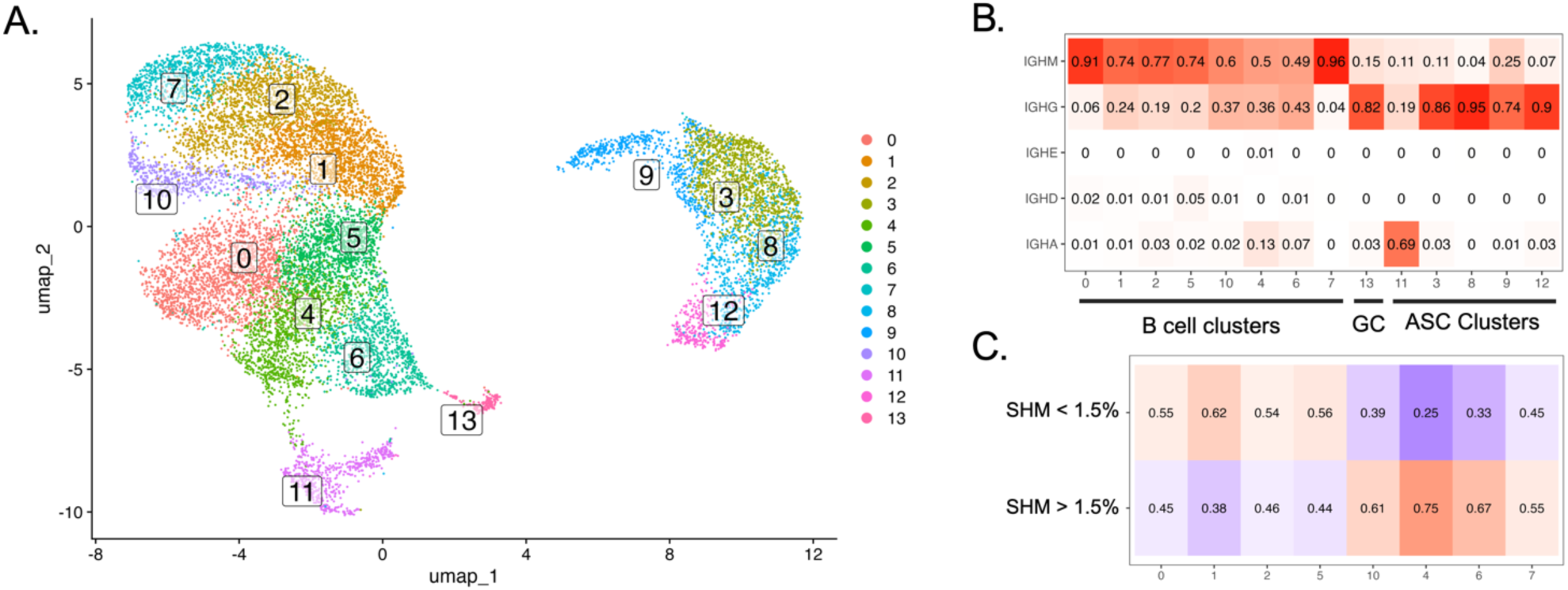
Enrichment of low mutation IGM/D expressing cells in distinct clusters. (A) UMAP of B lymphocyte lineage subclusters. (B) Heatmap of proportion of cells expressing each isotype within each subcluster; square color intensity and label based on the proportion. (C) Heatmap of proportion of IGHM expressing cells with low SHM (<1.5%) compared to high SHM (>1.5%) within the B cell subclusters; square color intensity and label based on the proportion. Created with BioRender.com

**Extended Data Figure 3.**
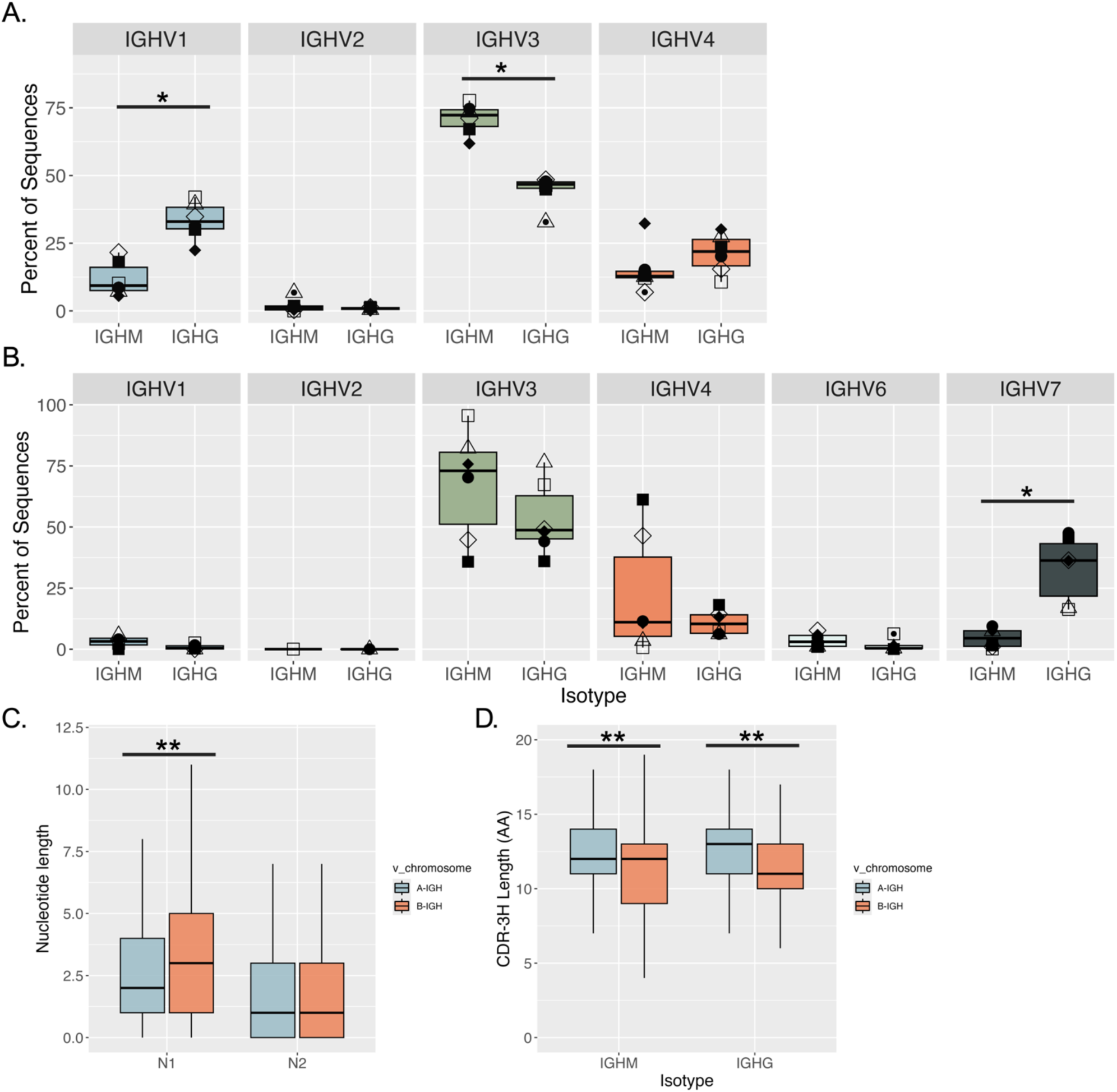
Analysis of Bulk IG repertoires. VH-family percentages for (A) A-IGH and (B) B-IGH within IGHM and IGHG from productive rearrangements found in bulk repertoire data where each point represents an individual bat (n = 6). (C) Box-and-whisker plot of length of V-D (N1 nucleotides) and D-J junctions (N2 nucleotides) for IGHM for A-IGH (blue) and B-IGH (orange). The horizontal line represents the median and whiskers span the range of the data excluding the outliers (C) Box-and-whisker plot of CDR-3H amino acid lengths including cysteine and tryptophan for IGHM and IGHG for A-IGH (blue) and B-IGH (orange). The horizontal line represents the median and whiskers span the range of the data excluding the outliers. *p < 0.005,**p < 0.000001, defined by clone wise unpaired Wilcoxon rank sum test. Graphs generated in R. Post-processing and figures created with BioRender.com

**Extended Data Figure 4.**
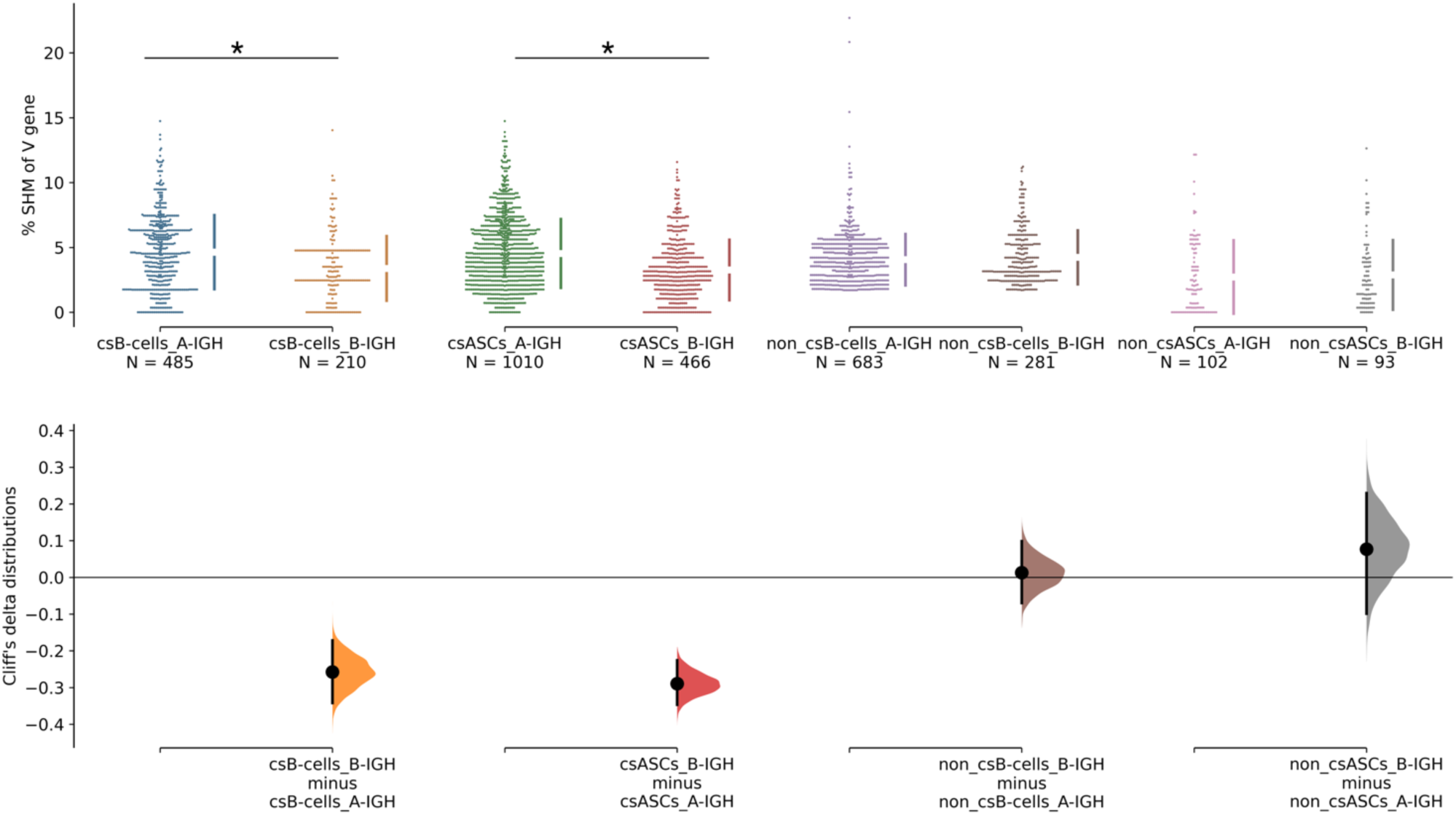
Permutation analysis of SHM. Top plot contains swarm plot of percent SHM of V gene sequences grouped by cell type comparing A-IGH (left) and B-IGH (right). Bottom plot is a main effect size plot non-paired Cliff’s delta with its 95% confidence interval for permutations of the non-parametric comparisons between A-IGH and B-IGH for each cell type. * p < 0.001

