## Supplemental Tables for "Genetically and Functionally Distinct Immunoglobulin Heavy Chain Locus Duplication in Bats"

**Supplementary Table 1. Annotations for A-IGH Constant genes**

| Locus | Designation | Exon | Genome Position [a] |
|---|---|---|---|
| A-IGH | A-IGHM | CH1 | 254914 - 255232 |
| | | CH2 | 255317 - 255652 |
| | | CH3 | 255914 - 256228 |
| | | CH4 | 256572 - 256904 |
| | | CH-S | 256905 - 256966 |
| | | TM1 | 259152 - 259267 |
| | | TM2 | 259353 - 259361 |
| A-IGH | A-IGHD | CH1 | 269606 - 269870 |
| | | H1 | 270195 - 270284 |
| | | H2 | 271683 - 271765 |
| | | CH3 | 271766 - 272099 |
| | | TM1 | 274229 - 274389 |
| | | TM2 | 274390 - 274395 |
| A-IGH | A-IGHG | CH1 | 296810 - 297095 |
| | | H1 | 297371 - 297415 |
| | | CH2 | 297507 - 297836 |
| | | CH3 | 297926 - 298246 |
| | | CH-S | 298247 - 298254 |
| | | TM1 | 299615 - 299745 |
| | | TM2 | 300352 - 300435 |
| A-IGH | A-IGHE | CH1 | 320077 - 320383 |
| | | CH2 | 320502 - 320822 |
| | | CH3 | 320907 - 321227 |
| | | CH4 | 321310 - 321636 |
| | | CH-S | 321637 - 321644 |
| | | TM1 | 324629 - 324730 |
| | | TM2 | 324851 – 324958 |
| A-IGH | A-IGHA | CH1 | 333612 - 333918 |
| | | H1 | 334117 - 334157 |
| | | CH2 | 334158 - 334464 |
| | | CH3 | 334653 - 334985 |
| | | CH-S | 334986 - 335047 |
| | | TM | 338349 – 338578 |

[a] Contig NC_072477.1

**Supplementary Table 2. Annotations for B-IGH Constant genes**

| Locus | Designation | Exon | Genome Position [b] |
|---|---|---|---|
| B-IGH | B-IGHM | CH1 | 1784216 - 1784534 |
| | | CH2 | 1784607 - 1784945 |
| | | CH3 | 1785177 - 1785491 |
| | | CH4 | 1785820 - 1786152 |
| | | CH-S | 1786153 - 1786214 |
| | | TM1 | 1788792 - 1788907 |
| | | TM2 | 1789001 - 1789009 |
| B-IGH | B-IGHD | CH1 | 1801850 - 1802113 |
| | | H1 | 1802396 - 1802482 |
| | | H2 | 1803897 - 1803979 |
| | | CH3 | 1803980 - 1804313 |
| | | TM1 | 1806140 - 1806288 |
| | | TM2 | 1806289 - 1806295 |
| B-IGH | B-IGHG | CH1 | 1818229 - 1818514 |
| | | H1 | 1818752 - 1818793 |
| | | CH2 | 1818888 - 1819217 |
| | | CH3 | 1819307 - 1819627 |
| | | CH-S | 1819628 - 1819635 |
| | | TM1 | 1820925 - 1821055 |
| | | TM2 | 1829396 - 1829479 |
| B-IGH | B-IGHE | CH1 | 1859123 - 1859429 |
| | | CH2 | 1859567 - 1859887 |
| | | CH3 | 1859954 - 1860292 |
| | | CH4 | 1860370 - 1860699 |
| | | CH-S | 1860700 - 1860707 |
| | | TM1 | 1864143 - 1864276 |
| | | TM2 | 1864623 - 1864730 |
| B-IGH | B-IGHA | CH1 | 1873005 - 1873308 |
| | | H1 | 1873478 - 1873500 |
| | | CH2 | 1873501 - 1873807 |
| | | CH3 | 1873998 - 1874330 |
| | | CH-S | 1874331 - 1874392 |
| | | TM | 1876670 - 1876899 |

[b] Contig NC_072496.1

**Supplementary Table 3. Nomenclature for IGHJ genes**

| Locus | Designation | Expressed | Genome Position |
|---|---|---|---|
| A-IGH | A-IGHJ1 | Y [c,d] | 247052 – 247105[a] |
| A-IGH | A-IGHJ2 | Y [c,d] | 247365 - 247414[a] |
| A-IGH | A-IGHJ3 | Y [c,d] | 247697 - 247744[a] |
| A-IGH | A-IGHJ4 | Y [c,d,e] | 248015 - 248062[a] |
| A-IGH | A-IGHJ5 | Y [c,d] | 248398 - 248448[a] |
| A-IGH | A-IGHJ6 | Y [c,d] | 248930 - 248983[a] |
| B-IGH | B-IGHJ1 | Y [c,d] | 1777158 - 1777211[b] |
| B-IGH | B-IGHJ2[ORF] | Y [c] | 1777808 - 1777855[b] |
| B-IGH | B-IGHJ3 | Y [c,d] | 1777808 - 1777855[b] |
| B-IGH | B-IGHJ4 | Y [c,d] | 1778083 - 1778130[b] |
| B-IGH | B-IGHJ5 | Y [c,d] | 1778466 - 1778516[b] |
| B-IGH | B-IGHJ6 | Y [c,d] | 1778884 - 1778937[b] |

[a] Contig NC_072477.1; [b] Contig NC_072496.1; [c] Functional rearranged in at least one individual in bulk IG repertoire; [d] Functional rearranged in $\geq$ 1 individual in single cell repertoire; [P] Pseudogene; [ORF] Open reading frame; [e] Putative allele found in $\geq$ 1 individual

**Supplementary Table 4. Nomenclature for IGHD genes**

| Locus | Designation | Length (nt) | Expressed [c,d] | Genome Position |
|---|---|---|---|---|
| A-IGH | A-IGHD1 | 31 | Y | 233577 - 233607[a] |
| A-IGH | A-IGHD2 | 22 | Y | 233913 - 233934[a] |
| A-IGH | A-IGHD3 | 19 | Y | 234857 - 234875[a] |
| A-IGH | A-IGHD4 | 20 | Y | 235089 - 235108[a] |
| A-IGH | A-IGHD5 | 16 | Y | 235511 - 235526[a] |
| A-IGH | A-IGHD6 | 10 | Y | 236216 - 236225[a] |
| A-IGH | A-IGHD7 | 19 | Y | 236580 - 236598[a] |
| A-IGH | A-IGHD8 | 17 | Y | 236812 - 236828[a] |
| A-IGH | A-IGHD9 | 16 | Y | 236984 - 236999[a] |
| A-IGH | A-IGHD10 | 10 | Y | 246699 - 246708[a] |
| B-IGH | B-IGHD1 | 33 | Y | 1763176 - 1763208[b] |
| B-IGH | B-IGHD2 | 13 | Y | 1763773 - 1763785[b] |
| B-IGH | B-IGHD3 | 15 | Y | 1763989 - 1764003[b] |
| B-IGH | B-IGHD4 | 16 | Y | 1766304 - 1766319[b] |
| B-IGH | B-IGHD5 | 10 | Y | 1767058 - 1767067[b] |
| B-IGH | B-IGHD6 | 19 | Y | 1767404 - 1767422[b] |
| B-IGH | B-IGHD7 | 18 | Y | 1767651 - 1767668[b] |
| B-IGH | B-IGHD8 | 16 | Y | 1767830 - 1767845[b] |
| B-IGH | B-IGHD9 | 10 | Y | 1776812 - 1776821[b] |

[a] Contig NC_072477.1; [b] Contig NC_072496.1; [c] Functional rearranged in $\geq$ 1 individual in bulk analysis; [d] Functional rearranged in $\geq$ 1 individual in single cell analysis [P] Pseudogene, [ORF] Open reading frame

**Supplementary Table 5. Nomenclature of A-IGH Variable Genes**

| Family | Locus | Proposed Designation | Expressed | Allele(s) | Human homolog | Genome Position[a] |
|---|---|---|---|---|---|---|
| 1 | A-IGH | A-IGHV1-1[P] | -- | -- | IGHV1-2 | 231604 - 229122 |
| 1 | A-IGH | A-IGHV1-5[P] | -- | -- | IGHV1-46 | 201160 - 196756 |
| 1 | A-IGH | A-IGHV1-12[P] | -- | -- | IGHV1-69 | 158537 - 153960 |
| 1 | A-IGH | A-IGHV1-13 | Y[b,c] | Y[d] | IGHV1-46 | 153490 - 148227 |
| 1 | A-IGH | A-IGHV1-14 | Y[b,c] | Y | IGHV1-46 | 147757 - 145163 |
| 1 | A-IGH | A-IGHV1-18 | Y[b,c] | -- | IGHV1-46 | 133495 - 129585 |
| 1 | A-IGH | A-IGHV1-20 | Y[b,c] | Y | IGHV1-46 | 126195 - 122974 |
| 1 | A-IGH | A-IGHV1-27 | Y[b,c] | Y | IGHV1-46 | 92354 - 92825 |
| 2 | A-IGH | A-IGHV2-3 | Y[b,c] | Y | IGHV3-30 | 212751 - 204865 |
| 2 | A-IGH | A-IGHV2-6 | Y[b,c] | Y | IGHV4-4 | 196285 - 190476 |
| 2 | A-IGH | A-IGHV3-9 | Y[b,c] | -- | IGHV3-48 | 180693 - 170120 |
| 2 | A-IGH | A-IGHV3-11 | Y[b,c] | Y | IGHV3-69-1 | 163267 - 159007 |
| 3 | A-IGH | A-IGHV3-2 | Y[b,c] | Y | IGHV3-21 | 228634 - 213222 |
| 3 | A-IGH | A-IGHV3-4 | Y[b,c] | Y | IGHV3-21 | 204382 - 201627 |
| 3 | A-IGH | A-IGHV3-7 | Y[b,c] | -- | IGHV3-11 | 189984 - 185742 |
| 3 | A-IGH | A-IGHV3-9 | Y[b,c] | Y | IGHV3-48 | 180693 – 170120 |
| 3 | A-IGH | A-IGHV3-15[ORF] | Y[b] | -- | IGHV3-48 | 144676 - 140485 |
| 3 | A-IGH | A-IGHV3-16[P] | -- | -- | pIGHV3-48 | 139991 - 137248 |
| 3 | A-IGH | A-IGHV3-17 | Y[b,c] | -- | IGHV3-69-1 | 136760 - 133965 |
| 3 | A-IGH | A-IGHV3-19[P] | -- | -- | IGHV3-48 | 129100 - 126665 |
| 3 | A-IGH | A-IGHV3-21 | Y[b,c] | Y | IGHV3-69-1 | 122480 - 118230 |
| 3 | A-IGH | A-IGHV3-23 | Y[b,c] | Y[d] | IGHV3-48 | 111260 - 107041 |
| 3 | A-IGH | A-IGHV3-24 | Y[a] | -- | IGHV3-48 | 103834 - 104323 |
| 3 | A-IGH | A-IGHV3-26 | Y[b,c] | Y | IGHV3-23 | 96569 - 97054 |
| 3 | A-IGH | A-IGHV3-29 | Y[b,c] | -- | IGHV3-21 | 79943 - 80431 |
| 3 | A-IGH | A-IGHV3-30 | Y[b,c] | Y | IGHV3-48 | 78023 - 78513 |
| 3 | A-IGH | A-IGHV3-31 | Y[b,c] | Y | IGHV3-69-1 | 74092 - 74577 |
| 3 | A-IGH | A-IGHV3-33 | Y[b,c] | Y | IGHV3-21 | 65288 - 65776 |
| 4 | A-IGH | A-IGHV4-8 | Y[b,c] | Y | IGHV4-30 | 185267 - 181181 |
| 4 | A-IGH | A-IGHV4-10 | Y[b,c] | Y | IGHV4-38 | 169649 - 163761 |
| 4 | A-IGH | A-IGHV4-22 | Y[b,c] | Y | IGHV4-28 | 117758 - 111749 |
| 4 | A-IGH | A-IGHV4-25 | Y[b,c] | -- | IGHV4-38 | 99145 - 99617 |
| 4 | A-IGH | A-IGHV4-28[P] | -- | -- | IGHV4-38 | 84996 - 85332 |
| 4 | A-IGH | A-IGHV4-32 | Y[b,c] | Y | IGHV4-38 | 70047 - 70523 |
| 4 | A-IGH | A-IGHV4-34 | Y[b,c] | Y | IGHV4-38 | 62260 - 62735 |

[a] Contig NC_072477.1, [b] Functional BCR in at least one individual in bulk analysis, [c] Functional BCR in at least one individual in single cell analysis, [P] Pseudogene, [ORF] open reading frame, [d] Putative allele found in $\geq$ 1 individual

**Supplementary Table 6. Nomenclature B-IGH variable genes**

| Family | Locus | Proposed designation | Expressed | Allele | Genome Position[a] |
|---|---|---|---|---|---|
| 1 | B-IGH | B-IGHV1-64 [P] | -- | -- | 16979486 - 16979961 |
| 1 | B-IGH | B-IGHV1-74 [P] | -- | -- | 17057215 - 17057678 |
| 1 | B-IGH | B-IGHV1-84 | Y[b,c] | Y | 17157598 - 17158077 |
| 1 | B-IGH | B-IGHV1-88 [P] | -- | -- | 17184987 - 17185458 |
| 1 | B-IGH | B-IGHV1-96 [P] | -- | -- | 17285813 - 17286389 |
| 1 | B-IGH | B-IGHV1-99 [ORF] | Y[b] | -- | 17301061 - 17301498 |
| 2 | B-IGH | B-IGHV2-51 [ORF] | N | -- | 16896960 - 16897380 |
| 2 | B-IGH | B-IGHV2-77 | Y[b] | -- | 17096111 - 17096580 |
| 3 | B-IGH | B-IGHV3-1 [P] | -- | -- | 16503414 - 16503816 |
| 3 | B-IGH | B- IGHV3-4 [ORF] | Y[b,c] | -- | 16527789 - 16528195 |
| 3 | B-IGH | B-IGHV3-7 [P] | -- | -- | 16556888 - 16557291 |
| 3 | B-IGH | B-IGHV3-8 | Y[b,c] | Y | 16560533 - 16560935 |
| 3 | B-IGH | B-IGHV3-10 | Y[b,c] | -- | 16570805 - 16571208 |
| 3 | B-IGH | B-IGHV3-12 [P] | -- | -- | 16582709 - 16583111 |
| 3 | B-IGH | B-IGHV3-14 [ORF] | Y[b,c] | Y | 16606837 - 16607244 |
| 3 | B-IGH | B-IGHV3-17 | Y[b,c] | -- | 16623445 - 16623848 |
| 3 | B-IGH | B-IGHV3-19 | Y[b] | -- | 16634141 - 16634544 |
| 3 | B-IGH | B-IGHV3-20 | Y[b,c] | Y[d] | 16638301 - 16638701 |
| 3 | B-IGH | B-IGHV3-22 | Y[b,c] | Y | 16650678 - 16651081 |
| 3 | B-IGH | B-IGHV3-24 | Y[b,c] | -- | 16665389 - 16665792 |
| 3 | B-IGH | B-IGHV3-26 [P] | -- | -- | 16681975 - 16682378 |
| 3 | B-IGH | B-IGHV3-27 | Y[b,c] | -- | 16685649 - 16686052 |
| 3 | B-IGH | B-IGHV3-28 | Y[b,c] | Y | 16696254 - 16696657 |
| 3 | B-IGH | B-IGHV3-29 | Y[b,c] | -- | 16700373 - 16700773 |
| 3 | B-IGH | B-IGHV3-30 | Y[b,c] | -- | 16716588 - 16716990 |
| 3 | B-IGH | B-IGHV3-33 [P] | -- | -- | 16733309 - 16733715 |
| 3 | B-IGH | B-IGHV3-36 [P] | -- | -- | 16750812 - 16751226 |
| 3 | B-IGH | B-IGHV3-37 | Y[b,c] | Y | 16764040 - 16764425 |
| 3 | B-IGH | B-IGHV3-41 | Y[b,c] | -- | 16810686 - 16811098 |
| 3 | B-IGH | B-IGHV3-42 [P] | -- | -- | 16831837 - 16832249 |
| 3 | B-IGH | B-IGHV3-43 [P] | -- | -- | 16833699 - 16833900 |
| 3 | B-IGH | B-IGHV3-46 | Y[b,c] | Y | 16850124 - 16850509 |
| 3 | B-IGH | B-IGHV3-48 [P] | -- | -- | 16879081 - 16879082 |
| 3 | B-IGH | B-IGHV3-49 | Y[b] | -- | 16885573 - 16885973 |
| 3 | B-IGH | B-IGHV3-50 [P] | -- | -- | 16894044 - 16894455 |
| 3 | B-IGH | B-IGHV3-55 [P] | -- | -- | 16927207 - 16927588 |
| 3 | B-IGH | B-IGHV3-56 | Y[b,c] | -- | 16935682 - 16936094 |
| 3 | B-IGH | B-IGHV3-58 | Y[b,c] | -- | 16948156 - 16948563 |
| 3 | B-IGH | B-IGHV3-59 [P] | -- | -- | 16951922 - 16952367 |
| 3 | B-IGH | B-IGHV3-61 | Y[b,c] | -- | 16964700 - 16965184 |
| 3 | B-IGH | B-IGHV3-62 | Y[b,c] | -- | 16968603 - 16969101 |
| 3 | B-IGH | B-IGHV3-67 [P] | -- | -- | 17018077 - 17018615 |
| 3 | B-IGH | B-IGHV3-68 [P] | -- | -- | 17028332 - 17028823 |
| 3 | B-IGH | B-IGHV3-69 | N | -- | 17030286 - 17030768 |
| 3 | B-IGH | B-IGHV3-70 | Y[c] | -- | 17034814 - 17035309 |
| 3 | B-IGH | B-IGHV3-72 | Y[b,c] | -- | 17048349 - 17048845 |
| 3 | B-IGH | B-IGHV3-75 [P] | -- | -- | 17066902 - 17067385 |
| 3 | B-IGH | B-IGHV3-76 | Y[b,c] | -- | 17079119 - 17079611 |
| 3 | B-IGH | B-IGHV3-79 | Y[b,c] | -- | 17125108 - 17125603 |
| 3 | B-IGH | B-IGHV3-80 [P] | -- | -- | 17132447 - 17132955 |
| 3 | B-IGH | B-IGHV3-81 [P] | -- | -- | 17145105 - 17145439 |

[a] Contig NC_072496.1; [b] Functional rearranged in ≥ 1 individual in bulk analysis; [c] Functional rearranged in ≥ 1 individual in single cell analysis; [P] Pseudogene; [ORF] Open reading frame; [d] Putative allele found in ≥ 1 individual

**Supplementary Table 6** (*continued*)

| Subgroup | Locus | Proposed designation | Expressed | Alleles | Genome Position[a] |
|---|---|---|---|---|---|
| 3 | B-IGH | B-IGHV3-82 [P] | -- | -- | 17147532 - 17147986 |
| 3 | B-IGH | B-IGHV3-83 [P] | -- | -- | 17154787 - 17155121 |
| 3 | B-IGH | B-IGHV3-86 [ORF] | Y[b,c] | -- | 17175826 - 17176321 |
| 3 | B-IGH | B-IGHV3-90 | Y[b,c] | -- | 17215824 - 17216311 |
| 3 | B-IGH | B-IGHV3-91 | Y[b,c] | -- | 17243422 - 17243916 |
| 3 | B-IGH | B-IGHV3-93 [P] | -- | -- | 17252274 - 17252571 |
| 3 | B-IGH | B-IGHV3-94 | Y[b,c] | -- | 17258745 - 17259242 |
| 3 | B-IGH | B-IGHV3-95 | Y[b,c] | Y | 17280061 - 17280555 |
| 4 | B-IGH | B-IGHV4-3 | Y[b,c] | Y | 16515583 - 16515972 |
| 4 | B-IGH | B-IGHV4-5 [P] | -- | -- | 16536112 - 16538064 |
| 4 | B-IGH | B-IGHV4-6 | Y[b,c] | Y | 16541640 - 16542028 |
| 4 | B-IGH | B-IGHV4-11 | Y[b,c] | -- | 16579921 - 16580313 |
| 4 | B-IGH | B-IGHV4-13 | Y[b] | -- | 16586700 - 16587088 |
| 4 | B-IGH | B-IGHV4-15 | Y[b,c] | -- | 16615433 - 16615822 |
| 4 | B-IGH | B-IGHV4-16 [P] | -- | -- | 16622277 - 16622278 |
| 4 | B-IGH | B-IGHV4-23 [P] | -- | -- | 16656563 - 16656998 |
| 4 | B-IGH | B-IGHV4-31 [P] | -- | -- | 16725032 - 16725033 |
| 4 | B-IGH | B-IGHV4-32 | Y[b] | -- | 16731799 - 16732191 |
| 4 | B-IGH | B-IGHV4-34 [P] | -- | -- | 16736829 - 16737222 |
| 4 | B-IGH | B-IGHV4-35 [P] | -- | -- | 16747513 - 16747607 |
| 4 | B-IGH | B-IGHV4-38 | N | -- | 16772220 - 16772606 |
| 4 | B-IGH | B-IGHV4-39 [P] | -- | -- | 16796802 - 16796918 |
| 4 | B-IGH | B-IGHV4-40 | Y[b,c] | Y | 16807104 - 16807498 |
| 4 | B-IGH | B-IGHV4-47 [P] | -- | -- | 16874859 - 16875153 |
| 4 | B-IGH | B-IGHV4-52 [P] | -- | -- | 16908253 - 16908635 |
| 4 | B-IGH | B-IGHV4-57 [P] | -- | -- | 16943608 - 16943970 |
| 4 | B-IGH | B-IGHV4-60 [P] | -- | -- | 16959989 - 16960423 |
| 4 | B-IGH | B-IGHV4-63 | Y[b,c] | -- | 16975868 - 16976344 |
| 4 | B-IGH | B-IGHV4-65 | Y[b,c] | -- | 16994204 - 16994681 |
| 4 | B-IGH | B-IGHV4-71[P] | -- | -- | 17036667 - 17037020 |
| 4 | B-IGH | B-IGHV4-73 | Y[b,c] | -- | 17053219 - 17053694 |
| 4 | B-IGH | B-IGHV4-78[P] | -- | -- | 17120700 - 17121032 |
| 4 | B-IGH | B-IGHV4-85 [P] | -- | -- | 17167544 - 17167884 |
| 4 | B-IGH | B-IGHV4-87 [P] | -- | -- | 17180661 - 17181023 |
| 4 | B-IGH | B-IGHV4-89 [P] | -- | -- | 17194181 - 17194492 |
| 4 | B-IGH | B-IGHV4-92 | Y[b] | -- | 17245898 - 17246375 |
| 4 | B-IGH | B-IGHV4-97 [P] | -- | -- | 17292871 - 17293407 |
| 4 | B-IGH | B-IGHV4-98 [P] | -- | -- | 17296958 - 17297686 |
| 5 | B-IGH | B-IGHV5-44 [P] | -- | -- | 16835449 - 16835450 |
| 5 | B-IGH | B-IGHV5-53 [P] | -- | -- | 16919228 - 16918916 |
| 6 | B-IGH | B-IGHV6-2 | Y[b,c] | -- | 16505787 - 16506187 |
| 6 | B-IGH | B-IGHV6-9 | N | -- | 16564103 - 16564503 |
| 6 | B-IGH | B-IGHV6-18 | Y[b,c] | Y | 16627033 - 16627433 |
| 6 | B-IGH | B-IGHV6-21 [P] | -- | -- | 16641247 - 16641342 |
| 6 | B-IGH | B-IGHV6-25 [P] | -- | -- | 16670778 - 16671178 |
| 7 | B-IGH | B-IGHV7-45 | Y[b,c] | -- | 16840998 - 16841387 |
| 7 | B-IGH | B-IGHV7-54 | Y[b,c] | Y | 16924172 - 16924561 |
| 7 | B-IGH | B-IGHV7-66 | Y[b,c] | -- | 17014838 - 17015314 |

[a] Contig NC_072496.1; [b] Functional rearranged in $\geq$ 1 individual in bulk analysis; [c] Functional rearranged in $\geq$ 1 individual in single cell analysis; [P] Pseudogene; [ORF] Open reading frame; [d] Putative allele found in $\geq$ 1 individual

**Supplementary Table 7. Nomenclature for IGLV genes**

| Family | Proposed designation | Orientation | Expressed | Allele | Genome Position[a] |
|---|---|---|---|---|---|
| 1 | IGLV1-1 | R | Y[b,c] | -- | 542686 - 524186 |
| 1 | IGLV1-2 | R | Y[b,c] | -- | 527862 - 527362 |
| 1 | IGLV1-3 | R | Y[b,c] | -- | 521727 - 521230 |
| 1 | IGLV1-4 | R | Y[b,c] | -- | 517896 - 517396 |
| 1 | IGLV1-5 | R | Y[b,c] | -- | 514165 - 513665 |
| 1 | IGLV1-6 | R | Y[b,c] | -- | 510669 - 510169 |
| 1 | IGLV1-7 | R | Y[b,c] | -- | 506960 - 506460 |
| 1 | IGLV1-8 | R | Y[b,c] | -- | 503693 - 503192 |
| 1 | IGLV1-9 | R | Y[b,c] | -- | 500099 - 499599 |
| 1 | IGLV1-58 | F | Y[b,c] | -- | 228608 - 229122 |
| 1 | IGLV1-85 [P] | F | -- | -- | 146790 - 147127 |
| 1 | IGLV1-90 | F | Y[b,c] | -- | 131365 - 131865 |
| 1 | IGLV1-96 [P] | F | -- | -- | 113044 - 113543 |
| 1 | IGLV1-102 [P] | F | Y[b,c] | -- | 94768 - 95272 |
| 1 | IGLV1-109 | F | Y[b,c] | Y[d] | 75840 - 76332 |
| 1 | IGLV1-113 [ORF] | F | N | -- | 62179 - 62760 |
| 1 | IGLV1-118 | F | Y[b,c] | -- | 46509 - 47009 |
| 1 | IGLV1-123 | F | Y[b,c] | -- | 31165 - 31668 |
| 2 | IGLV2-34 | F | Y[b,c] | -- | 334709 - 335212 |
| 2 | IGLV2-35 | F | Y[b,c] | -- | 332119 - 332623 |
| 2 | IGLV2-36 | F | Y[b,c] | -- | 329309 - 329812 |
| 2 | IGLV2-37 | F | Y[b,c] | -- | 326835 - 327329 |
| 2 | IGLV2-38 | F | Y[b,c] | -- | 324333 - 324835 |
| 2 | IGLV2-39 | F | Y[b,c] | -- | 321654 - 322157 |
| 2 | IGLV2-40 | F | Y[b,c] | -- | 319132 - 319635 |
| 2 | IGLV2-41 | F | Y[b,c] | -- | 316529 - 317033 |
| 2 | IGLV2-42 [P] | F | -- | -- | 313888 - 314392 |
| 2 | IGLV2-43 | F | Y[b,c] | -- | 308764 - 309267 |
| 2 | IGLV2-44 | F | Y[b,c] | -- | 302766 - 303268 |
| 2 | IGLV2-45 | F | Y[b,c] | -- | 300751 - 301263 |
| 2 | IGLV2-46 | F | Y[b,c] | Y[d] | 296116 - 296626 |
| 2 | IGLV2-47 | F | Y[b,c] | -- | 292268 - 292771 |
| 2 | IGLV2-48 | F | Y[b,c] | -- | 288407 - 288910 |
| 3 | IGLV3-10 | F | Y[b,c] | -- | 417299 - 418041 |
| 3 | IGLV3-11 | F | Y[b] | -- | 413465 - 414191 |
| 3 | IGLV3-12 | F | Y[b,c] | -- | 409398 - 409902 |
| 3 | IGLV3-14 | F | Y[b,c] | -- | 402436 - 402994 |
| 3 | IGLV3-16 | F | Y[b,c] | Y | 396020 - 396599 |
| 3 | IGLV3-18 | F | Y[b,c] | -- | 389289 - 390007 |
| 3 | IGLV3-19 | F | Y[b,c] | -- | 385460 - 385963 |
| 3 | IGLV3-21 | F | Y[b,c] | -- | 378575 - 379294 |
| 3 | IGLV3-22 [ORF] | F | Y[b,c] | -- | 375576 - 376071 |
| 3 | IGLV3-24 [P] | F | -- | -- | 369261 - 369983 |
| 3 | IGLV3-26 | F | Y[b,c] | -- | 362481 - 363201 |
| 3 | IGLV3-28 | F | Y[b,c] | -- | 355877 - 356459 |
| 3 | IGLV3-30 | F | Y[b,c] | -- | 349573 - 350017 |
| 3 | IGLV3-31 | F | Y[b,c] | -- | 345418 - 345923 |
| 3 | IGLV3-33 [ORF] | F | Y[b,c] | -- | 338573 - 339217 |
| 4 | IGLV4-13 | F | Y[b,c] | -- | 405385 - 405907 |
| 4 | IGLV4-15 | F | Y[b,c] | -- | 398776 - 399300 |
| 4 | IGLV4-17 | F | Y[b,c] | -- | 392204 - 392710 |

[a] Contig NC_072495.1; [b] Functional rearranged in ≥ 1individual in bulk analysis; [c] Functional rearranged in ≥ 1 individual in single cell analysis; [P] Pseudogene; [ORF] Open reading frame; [d] Putative allele found in ≥ 1 individual

**Supplementary Table 7.** (*continued*)

| Family | Proposed designation | Orientation | Expressed | Allele | Genome Position[a] |
|---|---|---|---|---|---|
| 4 | IGLV4-20 | F | Y[b,c] | -- | 381497 - 382020 |
| 4 | IGLV4-23 | F | Y[b,c] | -- | 372164 - 372687 |
| 4 | IGLV4-25 | F | Y[b,c] | -- | 365379 - 365892 |
| 4 | IGLV4-27 | F | Y[b,c] | -- | 358582 - 359095 |
| 4 | IGLV4-29 [P] | F | -- | -- | 352208 - 352732 |
| 4 | IGLV4-32 | F | Y[b,c] | -- | 341429 - 341954 |
| 5 | IGLV5-59 | F | Y[b,c] | -- | 224965 - 225477 |
| 5 | IGLV5-70 | F | Y[b,c] | -- | 188555 - 189068 |
| 5 | IGLV5-71 | F | Y[b,c] | -- | 185817 - 186319 |
| 5 | IGLV5-75 | F | Y[b,c] | -- | 173882 - 174405 |
| 5 | IGLV5-76 | F | Y[b,c] | -- | 170811 - 171324 |
| 5 | IGLV5-77 | F | Y[b,c] | -- | 167959 - 168480 |
| 5 | IGLV5-79 | F | Y[b,c] | -- | 163593 - 164106 |
| 5 | IGLV5-80 | F | Y[b,c] | -- | 160736 - 161244 |
| 5 | IGLV5-83 | F | Y[b,c] | -- | 152368 - 152889 |
| 5 | IGLV5-84 | F | Y[b,c] | -- | 150314 - 150819 |
| 5 | IGLV5-86 | F | Y[b,c] | -- | 144917 - 145425 |
| 5 | IGHV5-88 | F | Y[b,c] | -- | 137345 - 137866 |
| 5 | IGLV5-89 | F | Y[b,c] | -- | 134939 - 135452 |
| 5 | IGLV5-92 | F | Y[b,c] | -- | 126604 - 127125 |
| 5 | IGLV5-93 | F | Y[b,c] | -- | 123527 - 124040 |
| 5 | IGLV5-94 | F | Y[b,c] | -- | 119640 - 120156 |
| 5 | IGLV5-95 | F | Y[b,c] | -- | 115797 - 116310 |
| 5 | IGLV5-97 | F | Y[b,c] | Y | 109831 - 110336 |
| 5 | IGLV5-98 | F | Y[b,c] | -- | 107266 - 107787 |
| 5 | IGLV5-99 | F | Y[b] | -- | 104857 - 105370 |
| 5 | IGLV5-101 [P] | F | -- | -- | 97522 - 98034 |
| 5 | IGLV5-104 | F | Y[b,c] | -- | 89861 - 90369 |
| 5 | IGLV5-105 | F | Y[b,c] | -- | 86270 - 86783 |
| 5 | IGLV5-107 | F | Y[b,c] | -- | 80726 - 81239 |
| 5 | IGLV5-108 | F | Y[b,c] | -- | 78643 - 79167 |
| 5 | IGLV5-112 | F | Y[b,c] | -- | 65965 - 66467 |
| 5 | IGLV5-114 | F | Y[b,c] | -- | 60704 - 61206 |
| 5 | IGLV5-115 | F | Y[b,c] | -- | 55974 - 56498 |
| 5 | IGLV5-116 | F | Y[b,c] | -- | 53074 - 53587 |
| 5 | IGLV5-117 | F | Y[b,c] | -- | 50719 - 51249 |
| 5 | IGLV5-120 | F | Y[b,c] | -- | 40205 - 40707 |
| 5 | IGLV5-121 | F | Y[b,c] | -- | 37102 - 37617 |
| 5 | IGLV5-122 | F | Y[b,c] | -- | 33917 - 34430 |
| 5 | IGLV5-125 | F | Y[b] | -- | 26055 - 26563 |
| 5 | IGLV5-126 | F | Y[b,c] | -- | 22962 - 23483 |
| 5 | IGLV5-127 | F | Y[b,c] | -- | 20863 - 21401 |
| 6 | IGLV6-82 | F | Y[b,c] | -- | 155978 - 156497 |
| 6 | IGLV6-87 | F | Y[b,c] | -- | 140979 - 141490 |
| 7 | IGLV7-49 [ORF] | F | Y[b,c] | -- | 259071 - 259571 |
| 7 | IGLV7-50 [ORF] | F | Y[b,c] | -- | 256936 - 257427 |
| 7 | IGLV7-51 | F | Y[b,c] | -- | 253269 - 253741 |
| 7 | IGLV7-103 | F | Y[b,c] | -- | 91817 - 92289 |
| 7 | IGLV7-110 | F | Y[b,c] | -- | 72422 - 72891 |
| 7 | IGLV7-111 | F | Y[b,c] | -- | 69163 - 69635 |
| 7 | IGLV7-119 | F | Y[b] | -- | 43601 - 43944 |
| 7 | IGLV7-124 | F | Y[b,c] | -- | 28058 - 28486 |

[a] Contig NC_072495.1; [b] Functional rearranged in ≥ 1 individual in bulk analysis; [c] Functional rearranged in ≥ 1 individual in single cell analysis; [P] Pseudogene; [ORF] Open reading frame; [d] Putative allele found in ≥ 1 individual

**Supplementary Table 7.** (*continued)*

| Family | Proposed designation | Orientation | Expressed | Allele | Genome Position[a] |
|---|---|---|---|---|---|
| 8 | IGLV8-52 | F | Y[b,c] | -- | 249450 - 249945 |
| 8 | IGLV8-53 | F | Y[b,c] | -- | 244201 - 244692 |
| 8 | IGLV8-55 | F | Y[b,c] | Y | 238081 - 238572 |
| 8 | IGLV8-57 | F | Y[b,c] | -- | 231949 - 232440 |
| 8 | IGLV8-63 | F | Y[b,c] | -- | 212659 - 213150 |
| 8 | IGLV8-64 | F | Y[b,c] | -- | 208163 - 208654 |
| 8 | IGLV8-66 | F | Y[b,c] | -- | 202018 - 202509 |
| 8 | IGLV8-68 | F | Y[b,c] | -- | 195541 - 196032 |
| 8 | IGLV8-73 | F | Y[b,c] | -- | 179972 - 180464 |
| 8 | IGLV8-74 | F | Y[b,c] | -- | 177391 - 177886 |
| 10 | IGLV10-54 | F | Y[b,c] | -- | 240901 - 241415 |
| 10 | IGLV10-56 | F | Y[b,c] | -- | 234752 - 235266 |
| 10 | IGLV10-65 | F | Y[b,c] | -- | 204841 - 205355 |
| 10 | IGLV10-67 | F | Y[b,c] | -- | 198349 - 198863 |
| 10 | IGLV10-69 | F | Y[b,c] | -- | 192184 - 192698 |
| 11 | IGLV11-60 | F | Y[b,c] | -- | 222174 - 222676 |
| 11 | IGLV11-61 | F | Y[b,c] | Y | 218731 - 219213 |
| 11 | IGLV11-72 | F | Y[b,c] | -- | 182415 - 182892 |
| 11 | IGLV11-78 | F | Y[b,c] | -- | 166001 - 166522 |
| 11 | IGLV11-81 | F | Y[b,c] | -- | 158775 - 159296 |
| 11 | IGLV11-91 | F | Y[b,c] | -- | 128771 - 129248 |
| 11 | IGLV11-100 | F | Y[b,c] | -- | 101323 - 101838 |
| 11 | IGLV11-106 | F | Y[b,c] | -- | 84310 - 84831 |

[a] Contig NC_072495.1; [b] Functional rearranged in $\geq$ 1 individual in bulk analysis; [c] Functional rearranged in $\geq$ 1 individual in single cell analysis; [P] Pseudogene; [ORF] Open reading frame; [d] Putative allele found in $\geq$ 1 individual

**Supplementary Table 8. Nomenclature for IGLJ genes**

| Designation | Genome Position [a] |
|---|---|
| IGLJ1 | 428677 - 428714 |
| IGLJ2 | 433077 - 433114 |
| IGLJ3 | 436484 - 436521 |
| IGLJ4 | 439880 - 439917 |
| IGLJ5 | 443353 - 443390 |
| IGLJ6 | 446811 - 446848 |
| IGLJ7 | 450345 - 450382 |
| IGLJ8 | 453828 - 453865 |
| IGLJ9 | 457339 - 457376 |
| IGLJ10 | 460870 - 460907 |
| IGLJ11 | 464355 - 464392 |
| IGLJ12 | 467846 - 467883 |

[a] Contig NC_072495.1

\

**Supplementary Table 9. Nomenclature for IGLC genes**

| Designation | Genome Position [a] |
|---|---|
| IGLC1 | 430162 - 430480 |
| IGLC2 | 434420 - 434738 |
| IGLC3 | 437795 - 438113 |
| IGLC4 | 441223 - 441541 |
| IGLC5 | 444668 - 444987 |
| IGLC6 | 448188 - 448506 |
| IGLC7 | 451667 - 451985 |
| IGLC8 | 455178 - 455496 |
| IGLC9 | 458710 - 459028 |
| IGLC10 | 462192 - 462510 |
| IGLC11 | 465680 - 465998 |
| IGLC12 | 469099 - 469417 |

[a] Contig NC_072495.1
