## Supplemental Methods for "Genetically and Functionally Distinct Immunoglobulin Heavy Chain Locus Duplication in Bats"

### Animals

This study was carried out in accordance with recommendations set forth in the National Institutes of Health Guide for the Care and Use of Laboratory Animals[1]. Big brown bats (*Eptesicus fuscus*) used for this study were collected from a single colony in Calhoun, Georgia, United States. Collection of animals occurred under GA Department of Natural Resources permit #29-WSF-16-189. Capture, handling, and experimental procedures were performed in compliance with requirements of CDC Institutional Animal Care and Use Committee approved protocol 2809ELLBATC.

### *Eptesicus fuscus* immunoglobulin locus annotations

The *Eptesicus fuscus* DD_ASM_mEF_20220401 assembly and non-Ig gene annotations generated with the NCBI Eukaryotic Genome Annotation Pipeline from the NCBI database[2] were used to retrieve the contigs containing the Ig heavy and light sequences (or RPIA and EIF2AK3 in the case of the IGK locus). IgDetective[3] (v1.1.0) was used to identify contigs containing Ig heavy and light chain loci as well to predict putative V, J, and D genes. Additionally, variable, joining, and constant genes for heavy, lambda, and kappa loci from all species within the IMGT database[4] (v1.2.9) were mapped onto corresponding contigs using Geneious Prime 2021.1.1. Candidate gene segments were categorized as functional, open reading frame (ORF), and pseudogenes according to "Functionality" of IMGT[5]. Leader sequences were manually annotated. Recombination sequence signals (RSS) were identified using (https://www.itb.cnr.it/rss/analyze.html) and filtered based on previously described RIC score thresholds[6,7]. We classified V-gene units as pseudogenes or ORFs based on IMGT criteria[5].

Lastly, V gene annotations were validated and additional genes were identified by alignment of bulk IgM and IgG repertoires from 5'RACE (details below).

**AID hotspot density**

We used an interval spanning from the most 3' IGHJ gene to 5 kb after the TM domain of IGHA to search in both strands for the occurrence of the AID hotspot motif 5'-AGCT- 3'[8] using DNA-Pattern at RSA tools[9]. Raw counts were estimated in 1 kb non- overlapping windows and graph generated in R(v4.3.1)[10] and figure created using BioRender.com

**Comparative IGHV and IGLV phylogenetic analysis**

Functional, representative germline heavy and lambda variable genes from the major families for humans were collected from IMGT[10]. Multiple sequence alignment was performed on the *E. fuscus* and human IGHVs with MUSCLE and a nearest neighbor joining tree was generated with Jacks-Cantor using Geneious Prime 2021.1.1 (https://www.geneious.com). Final figure was generated using Fig.Tree (v1.4.4). For IGLV, the approach was identical.

**Comparative IGHM alignment**

Chromosomes and/or scaffolds containing immunoglobulin heavy chain constant regions were located and extracted using genome annotations and/or by applying BLAST[11] with IGHC genes from other species from publicly available bat genomes with a focus on vespertilionids with high quality genomes (**Methods Table 1)**. CH genes were initially annotated by mapping IGHC genes from other species to the regions and the IGHM region was aligned in Geneious Prime 2021.1.1 (https://www.geneious.com) and corrected manually. The final alignment spans from the beginning of the CH1 to ~1164 bp downstream of the end of the M domain. The phylogenetic hypothesis for the IGHM genes was generated using PhyML with a GTR+R substitution scheme which was the best supported model[12,13] and was run with 500 bootstrap replicates. The consensus phylogenetic hypothesis is displayed. Any nodes with less than 75% support were collapsed to a polytomy; the only unsupported nodes indicated an uncertainty in the placement of *Lasiurus cinereus* within the clade of "*Eptesicus*-like" (versus "*Myotis*-like") bats.

**Calculation of expected A-IGH:B-IGH ratio**

We calculated the expected ratio of A-IGH:B-IGH cells in naive populations under the assumption that recombination occurs in an ordered fashion starting at A-IGH as follows. Successful recombination assumed to occur 1/3 of the time such that productive A-IGH: 1/3 (allele #1A) + 2/3 * 1/3 (allele #2A) yields 0.56 for A-IGH and productive B-IGH: 2/3 * 2/3 *1/3 (allele #1B) + 2/3 * 2/3 * 2/3 *1/3 (allele #2B) yields 0.25. Therefore the A-IGH:B-IGH ratio is 0.56 / 0.25 or 2.24 for mature naive cells.

**A-IGH:B-IGH ratio**

Within each group (i.e. naive, non-class switched B cells, etc.), sequences were grouped by bat. Then the number of cells with BCRs from A-IGH plus a pseudo count of one was divided by the number of cells with BCRs from B-IGH plus a pseudo count of one. Frequency of A-IGH vs B-IGH for cumulative data were compared with a chi-square test.

**Bulk immunoglobulin repertoire methods**

RNA was isolated from flash frozen *Eptesicus fuscus* splenic tissue using Zymogen quick DNA/RNA mini kit (cat#D7001) and 5'RACE cDNA was generated using SMRTR 3'/5'RACE kit (Takara, cat#634858). Nested constant region primers for IGHM, IGHG, and IGLCs (sequences available pre-publication upon request) were generated from known sequences and chromosome level assembly annotations generated above. A touchdown PCR protocol was used for the first round PCR as follows: 1) 5 cycles of 94°C for 30s and 72°C for 3 minutes, 2) 5 cycles of 94°C for 30s, 70°C for 30s and 72°C for 3 minutes, and 3) 25 cycles of 94°C for 30s,

68°C for 30s, and 72°C for 3 minutes. The PCR product from round 1 was diluted in TE buffer and used as a template for a second round of three step PCR with an initial denaturation of 98°C for 30s, then 25 cycles of 98°C for 10s, 60°C for 30s and 72°C for 30s and final extension of 5 minutes. A dual sided SPRIselect (Beckman Coulter, cat#B23317) bead size selection was performed on PCR products. Libraries were barcoded using NEBNext DNA library kit (New England Biolabs, #E7645S and #E7370L). Libraries were pooled and sequenced using a MiSeq Reagent Kit v3 600 cycle (Illumina cat#MS-102-3003) on an Illumina MiSeq instrument. Reads were merged and trimmed using FLASH and aligned to the genome using Geneious Prime 2021.1.1 to validate annotations and identify additional genes. Data were then processed using MiXCR[14] (v4.6.0) with a custom reference annotation of BCRs and used to generate inferred allelic germline reference sets for each individual animal to be used below for single cell analysis.

**Single cell transcriptomic data generation**

Single cell suspensions were generated from a portion of fresh whole spleens and cryopreserved in complete media with 10% DMSO. Cells were thawed, washed once, counted, and resuspended in a 1x PBS + 0.2% BSA. Approximately 20,000 cells from a single bat were loaded per 10x lane. The Chromium GEM Single cell 5' kit v2 (10x Genomics PN-1000374) was used and protocol Rev D was followed for the preparation of cDNA and gene expression library generation.

For BCR enrichment libraries, nested constant region primers were generated from known sequences and chromosome level assembly annotations (sequences available pre-publication upon request). Custom outer constant region reverse primers were pooled and added (0.2 uM final) to 50 uL Amp Mix (10x Genomics, cat#2000047), forward primer (IDT based on

10x Genomics published sequence, 0.2uM), 2uL cDNA, and water. Thermocycler with lid set to 105°C was cycled as follows: 1) 98°C for 45s; 2) 98°C for 20s, 55°C for 30s, 72°C for 1 minute for 8 to 12 cycles; 3) 72°C for 1 min; and 4) hold at 4°C. After double sided bead selection, 35uL of sample was added to pooled custom inner constant region reverse primers (0.2 uM final), 50 uL Amp Mix, forward primer (IDT based on 10x Genomics published sequence, 0.2uM), and water. Thermocycler with lid set to 105°C was cycled as follows: 1) 98°C for 45s; 2) 98°C for 20s, 62°C for 30s, 72°C for 1 minute for 8 to 12 cycles; 3) 72°C for 1 min; and 4) hold at 4°C. The final amplified products were processed, libraries prepared and barcoded as detailed in the 10x Genomics protocol Rev D.

After library prep, quality control was performed using a bioanalyzer (Agilent 2100 Bioanalyzer, Agilent technologies) and quantification, pooling and dilution of libraries to 1.5nM was performed using KAPA library quantification kit (Roche, cat#07960140001). Gene expression libraries were pooled and sequenced using a Nova-seq SP 200 cycle kit (Illumina, San Diego, CA; cat#20040719) on an Illumina NovaSeq instrument. Gene expression and BCR libraries were pooled and sequenced using a Nova-seq S2, 200 cycle kit (cat#20028315) on an Illumina NovaSeq instrument. Raw data were deposited at [*Database*] and are accessible via *accession #*. The code used in this study can be obtained upon request.

**Single cell transcriptomic data analysis**

Custom cellranger reference was generated using the DD_ASM_mEF_20220401 assembly and annotations available on NCBI. Annotations for IG genes were removed and replaced with custom annotations. Single cell gene expression (GEX) data was processed using Cell Ranger (10x Genomics, v.8.0.0) followed by Seurat[15] (v5.0.1). Quality control measures were first applied to filter out cells with an unusually high or low number of detected genes, indicative of potential cell stress or death. Normalization of the data was performed using the NormalizeData function to mitigate the influence of cell-specific biases. The FindVariableFeatures function identified highly variable genes across the dataset, which were used for downstream analysis. Prior to initial clustering, IG genes were removed from the highly variable list. Datasets were integrated using CCA and layers were joined. Clustering was repeated and the dead cell cluster was removed. Principal component analysis (PCA) was conducted using the RunPCA function on the scaled data to reduce dimensionality. The significant principal components were selected based on the Elbow plot, guiding the selection of dimensions for clustering. Cell clusters were identified using the FindClusters function. The RunUMAP function was then employed to visualize the cells in a two-dimensional uniform manifold approximation and projection (UMAP) plot. Differential expression analysis between identified cell clusters was performed using the FindAllMarkers function with a Wilcoxon rank sum test. Marker genes were used to annotate clusters based on known cell type-specific expression profiles. Low-quality cells were discriminated by distinct clustering, expression of only mitochondrial and ribosomal genes, and lack of other phenotypic gene expression.

To identify B lymphocyte subsets, B lymphocyte lineage clusters and dividing cells were re-clustered; light chain genes were removed from the variable gene set but isotype genes were included. Cells were clustered and non-B cells and doublets formed distinct clusters and were removed. Remaining cells were re-clustered to yield the final subsets.

**Single cell BCR data analysis**

Paired-end reads generated by Single-Cell BCR sequencing were assembled into contigs representing full-length V(D)J recombinations for IGH and CDR-3 for IGL using MiXCR[14] (v4.6.0). Individual germline references containing putative alleles generated from bulk IG data (above) were used to process the data for each individual. The number of nucleotide changes per sequence was determined and normalized to the length of the V gene region to calculate the mutation frequency. For cell barcodes with multiple IGH contigs, a dominant contig was identified if 1) one chain had more than 5 times the reads of the other or 2) 3x more UMIs and the isotype for the dominant chain agreed with the transcriptomic expression for that cell. For IGH assemblies without an isotype, gene expression data from the single cell transcriptomes were leveraged to identify the isotype. An isotype was called in cases where only a single constant gene has non-zero count or if the same constant gene from both loci have non-zero counts (e.g., both A-IGHG and B-IGHG). Assemblies were then filtered to only include those which were identified as B lineage cells in the transcriptomic analysis.

**Selection Pressure Analysis**

Single cell IGH BCR sequences with germline reference were exported in the IMGT format using MiXCR (v4.6.0)[14]. Sequences were filtered for gaps and then Shazam[16–20] (v1.2.0) BASELINe was applied for analysis of selection pressure**.** Visualizations were generated in R using the SHazaM (v1.2.0) and then processed with Biorender.com.

**Statistical Analysis**

All statistical analyses were conducted in R (**Methods Table 2**). All comparisons are clone-wise for bulk IG data where a clone is defined as a unique sequence or cell-wise for single cell data unless otherwise stated. Non-paired Wilcoxon rank-sum tests were used to conduct pairwise comparisons. Multiple comparisons were conducted using Kruskal-Wallis rank sum tests followed by Dunn’s multiple test with Benjamini-Yekutieli for FDR correction. Permutation tests were performed in python with the DABEST[21] package using 5,000 permutations of unpaired Cliff’s delta between the groups. The 95% confidence interval was bias-corrected and accelerated. Statistical significance markers were added to graphs using ggpubr[22] package or manually added during figure generation with Biorender.com.

**Methods Table 1. Vespertilionid genomes used for comparative IGHM analysis**

| Species | Assembly (GenBank ID) |
|---|---|
| *Phyllostomus discolor* | GCA_004126475.3 |
| *Miniopterus natalensis* | GCA_001595765.1 |
| *Myotis brandtii* | GCA_000412655.1 |
| *Myotis yumanensis* | GCA_028538775.1 |
| *Corynorhinus townsendii* | GCA_026230045.1 |
| *Plecotus auritus* | GCA_963455305.1 |
| *Eptesicus fuscus* | GCA_027574615.1 |
| *Eptesicus nilssonii* | GCA_951640355.1 |
| *Ia io* | GCA_025583905.1 |
| *Pipistrellus pygmaeus* | GCA_949987765.1 |
| *Pipistrellus pipistrellus* | GCA_903992545.1 |
| *Pipistrellus kuhlii* | GCA_014108245.1 |
| *Lasiurus cinereus* | GCA_011751065.1 |

**Methods Table 2. Additional Software Packages**

| Package | Version |
|---|---|
| base[22] | 4.3.1 |
| Biostrings[23] | 2.70.3 |
| data.table[24] | 1.15.4 |
| dunn.test[25] | 1.3.6 |
| edgeR[26–28] | 4.0.16 |
| ggbeeswarm[29] | 0.7.2 |
| ggpubr[30] | 0.6.0 |
| ggrepel[31] | 0.9.5 |
| patchwork[32] | 1.2.0 |
| pheatmap[33] | 1.0.12 |
| reshape2[34] | 1.4.4 |
| Rmarkdown[35–37] | 2.26 |
| scales[38] | 1.3.0 |
| seqinr[39] | 4.2.36 |
| Seurat[40–43] | 5.0.3 |
| shazam[16,17,19] | 1.2.0 |
| stringdist[44] | 0.9.12 |
| tidyverse[21] | 2.0.0 |

**Methods References**


1. National Research Council, Division on Earth and Life Studies, Institute for Laboratory Animal Research & Committee for the Update of the Guide for the Care and Use of Laboratory Animals. *Guide for the Care and Use of Laboratory Animals: Eighth Edition*. (National Academies Press, 2010).
2. Thibaud-Nissen, F. *et al.* P8008 The NCBI Eukaryotic Genome Annotation Pipeline. *J. Anim. Sci.* **94**, 184–184 (2016).
3. Sirupurapu, V., Safonova, Y. & Pevzner, P. A. Gene prediction in the immunoglobulin loci. *Genome Res.* **32**, 1152–1169 (2022).
4. Giudicelli, V. *et al.* IMGT/LIGM-DB, the IMGT comprehensive database of immunoglobulin and T cell receptor nucleotide sequences. *Nucleic Acids Res.* **34**, D781-4 (2006).
5. Lefranc, M. P. Immunoglobulin and T Cell Receptor Genes: IMGT(®) and the Birth and Rise of Immunoinformatics. *Front. Immunol.* **5**, 22 (2014).
6. Cowell, L. G., Davila, M., Kepler, T. B. & Kelsoe, G. Identification and utilization of arbitrary correlations in models of recombination signal sequences. *Genome Biol.* **3**, RESEARCH0072 (2002).
7. Merelli, I. *et al.* RSSsite: a reference database and prediction tool for the identification of cryptic Recombination Signal Sequences in human and murine genomes. *Nucleic Acids Res.* **38**, W262-7 (2010).
8. Xu, Z., Zan, H., Pone, E. J., Mai, T. & Casali, P. Immunoglobulin class-switch DNA recombination: induction, targeting and beyond. *Nat. Rev. Immunol.* **12**, 517–531 (2012).
9. Medina-Rivera, A. *et al.* RSAT 2015: Regulatory Sequence Analysis Tools. *Nucleic Acids Res.* **43**, W50-6 (2015).
10. R Core Team. R: A Language and Environment for Statistical Computing. *https://www.R-project.org/*.

27. *EdgeR 4.0: Powerful Differential Analysis of Sequencing Data with Expanded Functionality and Improved Support for Small Counts and Larger Datasets*.
